# Year-to-year crop shifts promote weed diversity in tropical permanent rainfed cultivation

**DOI:** 10.1101/2020.03.12.988469

**Authors:** Margot Neyret, Anneke de Rouw, Nathalie Colbach, Henri Robain, Bounsamay Soulileuth, Christian Valentin

## Abstract

In the past decades, the expansion and modernisation of agriculture in the mountainous areas of Southeast Asia has had severe impacts on biodiversity, as the once species-rich forests were turned into homogeneous fields receiving ample external inputs. A common feature of permanent cropping with annual crops is the frequent change of crop choice, depending on market opportunities or other motives. However, the precise effect of crop shifts on weeds in tropical areas is largely unknown. In this study, we investigated the short-term effect of crop sequences on the diversity of weed communities in smallholder fields in Northern Thailand. Crop choices were upland rice, maize, fallow and young tree plantations with or without intercrop. We counted the number of crop shifts and the number of crops involved during a 3-years period preceding weed sampling. We showed that the number of crop shifts did not affect weed density and biomass. However, herbaceous species number and diversity (measured as Shannon index) increased by 36% and 46% respectively, while herbaceous species dominance decreased by 38%, in fields with yearly crop shifts compared to fields with no shifts in the previous three years. The effect of a particular crop on diversity, or the effect of intercropping with young trees, was weaker. It was likely due to the more variable resources (especially light) in fields with two crop shifts, allowing species with different niches to co-exist. Crop type and frequent crop shifts did not affect shrub and tree species number, diversity or dominance. Some species were strongly associated with fields with no crop shift in the sequence (e.g. the tree *Antidesma velutinosum*) or to fields with two crop shifts in the sequence (e.g. the herb *Centella asiatica*, the C_4_ grass *Digitaria radicosa*). Overall, this study showed that in this agronomical system, maintaining yearly crop shifts does not significantly affect weed abundance, but supports in-field plant species diversity, which is likely to impact the services provisioned by tropical mountainous agro-ecosystems

**Highlights:**

- Frequent crop shifts in a crop sequence increased weed richness and diversity.
- Crop shifts had a stronger effect on weed richness and diversity than the current crop.
- The number of crop shifts did not affect weed biomass and density.

**Graphical abstract:** 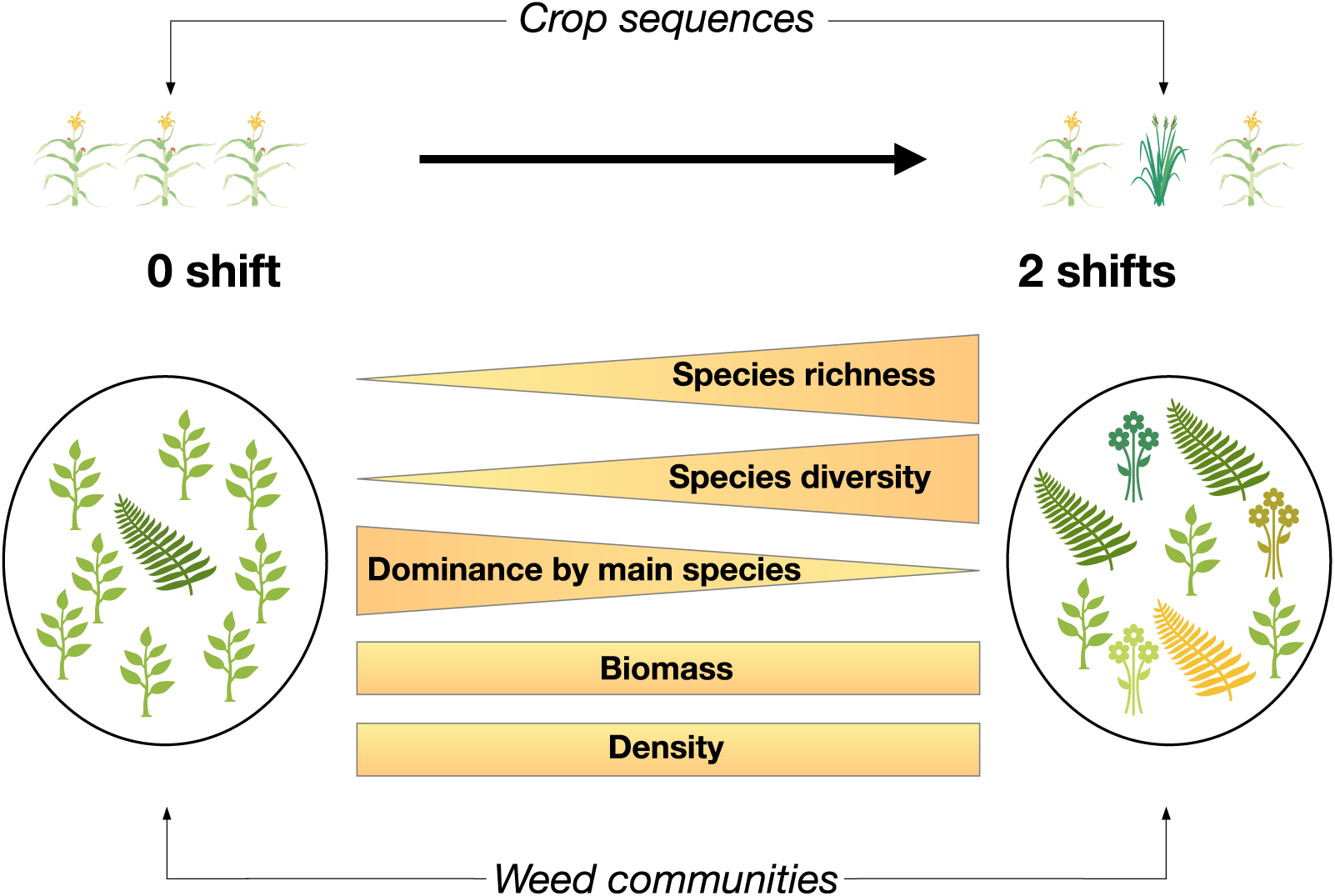

## 1. Introduction

The growing use of pesticides and fertilisers and the decrease of crop diversity associated with the modernisation of agriculture have led to a worldwide decrease of in-field biodiversity, soil quality and organic matter content, and an acceleration of surface water eutrophication (Tilman, 2001; Tscharntke et al., 2005). In temperate areas, agricultural intensification has been shown to strongly affect weed communities, by favouring species that are more competitive and mimic the main crop (Fried et al., 2010, 2012, 2015) and overall by decreasing in-field weed diversity and abundance (Squire et al., 2000; Hyvönen and Salonen, 2002; Baessler and Klotz, 2006; Fried et al., 2009; Hyvönen et al., 2011).

Yet, multiple ecosystem services depend on the maintenance of weed diversity (Matson et al., 1997; Tscharntke et al., 2005). Diverse plant communities are indicative of the wider sustainability and resistance to stress of the cropping system (Chen et al., 2004; Storkey and Neve, 2018). They provide food sources for biodiversity at higher trophic levels during extended periods (Marshall et al., 2003; Petit et al., 2011; Edesi et al., 2012) and promote large and rich populations of pollinators necessary to the cultivation of arthropod-pollinated crops (Bàrberi et al., 2010). Plant cover is also a major element of soil conservation by favouring infiltration, enhancing soil structure and organic carbon stocks (Durán Zuazo and Pleguezuelo, 2008). Contrasted plant traits have been shown to protect soil in various ways: the root density and the proportion of fine roots limit surface erosion (Burylo et al., 2012a, 2012b) while deeper roots stabilise slopes (Stokes et al., 2008). As a result, diverse weed communities enhance erosion mitigation: diverse root growth forms, for instance, have been shown to increase soil protection (Beierkuhnlein and Jentsch, 2004). Weeds also provide other key services such as pest control (Crowder and Jabbour, 2014), water filtration or nutrient cycling (Gholamhoseini et al., 2013a, 2013b; Yagioka et al., 2015), which are severely threatened by current agricultural changes.

From the 1970s onward, population growth and rapid economic development led to the expansion of arable lands to the detriment of natural and semi-natural areas and to a quick increase in the use of herbicides and fertilisers (Rerkasem and Rerkasem, 1995; Rigg et al., 2012). The development of cash crops was associated with the disruption of traditionally diverse cropping systems (Rerkasem et al., 2009). In Southeast Asia, one of the hotspots of biodiversity worldwide, these changes have occurred at an unprecedented scale (Fox et al., 2014). A better understanding of the relationships between these changes in cropping practices and non-cultivated biodiversity is particularly critical in mountainous areas, where biodiversity has been strongly affected by the recent modernisation of farming practices (Rerkasem et al., 2009).

Crop rotation is the practice of growing different crops on the same land from year to year following a more or less fixed cycle. Besides their well-known positive effects on soil health and fertility (Watson et al., 2002), erosion mitigation (Morgan, 2005), and crop disease risk reduction (Colbach et al., 1994), crop rotations are also an important method of weed control, as they prevent the build-up of aggressive weed communities linked to cultivating the same crop over and over again (Liebman and Dyck, 1993, Radosevich et al., 1997, Doucet et al., 1999; Squire et al., 2000 ; Adler et al., 2006; Ulber et al., 2009; Allan et al., 2014; Gaba et al., 2014). Most of these studies investigated the effect of well-defined rotations (Cardina et al., 2002; Nikolic et al., 2018; Shahzad et al., 2016) such as the common maize-soybean-wheat (Doucet et al., 1999) on weed communities and crop yield. A few studies have also proposed various metrics to define crop rotations, such as the proportion of a focus crop in the rotation. But the idea that crop rotations follow a fixed pattern can be questioned (von Redwitz and Gerowitt, 2018). In Southeast Asia, the rainfed cultivation such as the system we studied does not follow predetermined patterns, but rather results from year-to-year decisions based on the current meteorological or socio-economic context. There is an element of randomness that may be linked to the choices of individual farmers. The effect of these crop sequences on weed communities is unknown. It is urgent to investigate the weed diversity under any crop sequence, not only under the sequence of the rather standard and well-known crop rotations.

In this study, we determine the effect of crop sequences on the richness, diversity, biomass and density of weed communities in fifteen permanently cropped smallholder fields in mountainous Northern Thailand. We characterised the sequences using the number of crop shifts, and the number of crop types. We also aimed at identifying species that are indicators of fields with frequent crop shifts. We conducted weed sampling in the rainy season to collect additional information on agricultural practices and crop stand and in the dry season to collect information on the effect on weeds of post-harvest conditions such as water availability. We hypothesised that fields with frequent shifts would have more diverse and less abundant weed communities, especially in the dry season when the effect of the most recent crop is weaker, while weed biomass would be overall higher in the rainy season due to higher water availability.

## 2. Material and Methods

### 2.1. Study site

The study site was located in Huai Lang, Chiang Rai province, northern Thailand. The site is part of a wider project investigating the soil-water effects of land use transitions. The fields were located in and around two catchments, respectively dominated by annual cultivation or rubber tree plantations. An automatic weather station (Campbell BWS200) has been installed on-site since March 2015. The site is characterised by 1300 ± 200 mm of annual rainfall, mostly falling during the rainy season (May to November). Mean annual temperature is 24.5 ± 0.4C, with daily temperatures ranging from 4.8 to 42.5 C. A detailed soil mapping showed that soils are Haplustalfs (Alfisols) and belong to three soil series (Tha li, Wang Saphung, and Muak Lek, based on Jumpa (2012)), mostly differentiated on depth and slope criteria. The soils were otherwise rather uniform, well drained and with clay to clay-loam texture.

Paddies (wet rice), maize fields and settlements occupied most of the flatlands but we focused our study the rain-fed fields of the hillslopes, which have typically steep slopes ranging from 27 % to 54 % (median 40 %). The size of the study fields ranged from 0.64 ha to 2.6 ha (median 1.6 ha). On the hillslopes, maize (cash crop) and upland rice (subsistence crop) were grown in monoculture or as an intercrop between rows of young trees (most often immature rubber trees). Farmers prepared their fields between April and June and, with a few exceptions, they burnt crop residues before seeding. In a given field, maize was grown for one, two or three consecutive years and rice only for one or two years. Upland rice was planted at the beginning of the rainy season (late May-June) whereas the planting period of maize was more flexible. Indeed, being a crop with a short growing season, maize could be sown later in the rainy season (from May and up to July) and benefited from the long growing season associated with the bimodality of the climate. Maize and rice were harvested during October and November, respectively. The steep slopes did not permit ploughing and the soil was mostly left undisturbed, except for occasional manual surface hoeing. The upland rice varieties in the study area were long-cycle landraces (i.e. locally adapted varieties), typically tall varieties with drooping leaves providing dense shadow. In contrast, the maize varieties were modern, short-cycle and herbicide-resistant improved varieties grown as a cash crop for animal feed. Rice was planted in hills with an average density of 130 000 hill/ha, which was common for landraces in the area. Maize was sown in densities of 31 000 hill/ha, with two plants per hill, which was in the low range of typical sowing densities (20 000 hill/ha to 80 000 hill/ha). In both maize and rice glyphosate was the most common herbicide, applied with rates ranging from 0.7 L/ha to 25 L/ha (at 480 g/L) and often in combination with other herbicides such as paraquat or atrazine (Neyret et al., 2018). These values, displaying a surprisingly large range, were reported by farmers but could not be checked in the field. Herbicides were sprayed before or just after sowing in rice (May - June), and before sowing (April – May) and sometimes after emergence (July) in maize. Most farmers fertilised their field by applying 13 kg/ha to 130 kg/ha of fertiliser: usually urea (46-00) once a year, more rarely twice a year or in combination with ammonium sulphate (21-00) or NPK fertilizer (15-15-15).

Rubber (*Hevea brasiliensis*) is an important cash crop in the study area. The oldest plantations were planted at the beginning of the 2000s, and the expansion of rubber trees is ongoing. Saplings of rubber trees are planted on the slopes (approx. 500-550 trees/ha) and commonly intercropped with maize or upland rice during 4 – 5 years when young allowing farmers to improve their income. Intercropping of maize and upland rice with longan fruit trees (*Dimocarpus longan*) is also common. Longan is planted in low densities (approx. 300 trees/ha).

### 2.2. Sampling protocol

In March 2016, we selected 15 fields (five maize fields, five upland rice fields, five young rubber trees plantations with maize intercrop). We chose field locations in order to avoid spatial clustering of similar crops. This was done by selecting distinct sectors with at least 2-3 different crop types within a few hundred meters each. In each field, we delimited a 100-m^2^ area in a section representative of the whole field (i.e. avoiding field edges and large terrain irregularities). We then divided this area into a regular grid of ten by ten 1-m^2^ subplots and randomly drew 5 numbers between 1 and 100, which determined the position of five 1-m^2^ subplots in the grid. We conducted complete botanical inventories (e.g. individual plant counts and identification) in these subplots. We also collected the aboveground biomass of all living herbaceous weeds, shrubs and trees (thus excluding crop biomass); it was stored in paper bags and dried at 50 °C for 48 h before weighting. Biomass measurements were then averaged for each field. As resprouting trees and shrubs were less abundant than herbaceous species, we also counted and identified all trees and shrubs in the 100-m^2^ areas. Both herbaceous and shrub/tree densities exclude crops, and were then converted to the average number of plants per square meter for each 100-m^2^ area.

In total, this protocol was maintained in the same fields (but in different subplots) during three years. Sampling was conducted three times in the dry season (in March 2016, 2017 and 2018, before field preparation); and twice in the rainy season (October 2016 and November 2017, just after the harvest). Sometimes the crop was not fully mature and we sampled fields shortly before harvest - and thus did not collect data on yield. In the rainy season soil moisture was on average 2 to 3 times higher than in the dry season (Neyret, 2019). Samplings were conducted at least two (rainy season) or seven (dry season) months after the last herbicide application.

### 2.3. Diversity indices

We used multiple descriptors of weed communities that provided complementary information (Table 1). Plant biomass and density provided information on the potential aggressiveness of weed communities towards the crop. The number of weed species is a simple measure of plant richness, which we complemented by diversity and dominance indices – describing respectively the diversity of the whole community (Shannon index, noted H’) and the strength of the dominance by the main species (Berger-Parker index, noted D). Diversity and dominance are important indicators of the resistance and the resilience of an ecosystem, as more diverse communities are likely to be more stable and resilient (McCann, 2000).

**Table 1.**
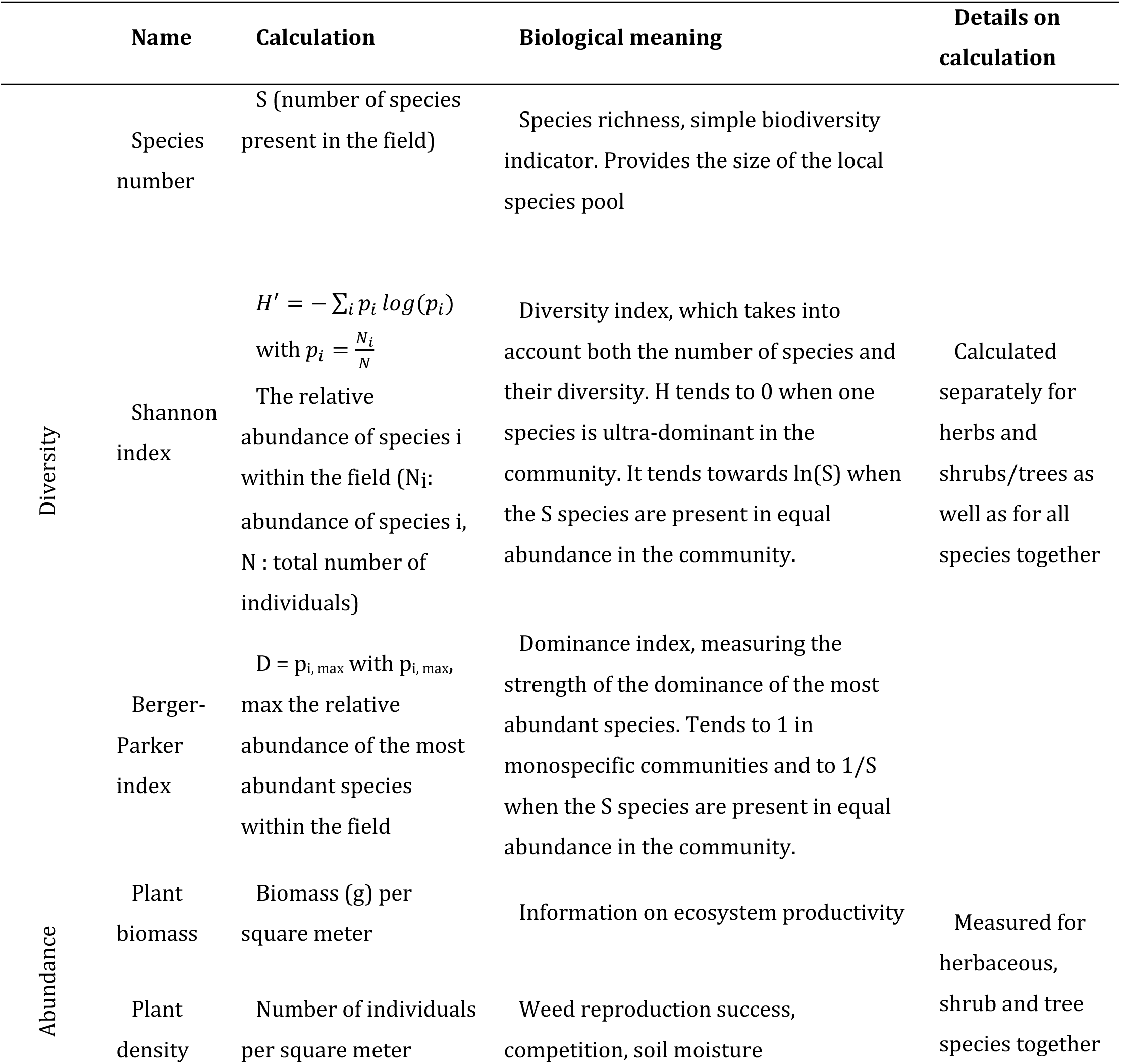
Selected descriptors of plant communities.

We ranked the species according to a Relative Importance index RI (e.g. Cardina et al., 2002), adapted from the Importance Value Index (which also takes into account biomass or basal area) to distinguish between the most common, intermediate and rare species.

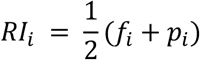

RI ranged from 0 to 1 and was calculated for each species i as the average of its relative frequency (fi, number of plots where species i was found divided by the total number of plots) and relative abundance (pi, number of individuals in species i divided by the total number of individuals).

### 2.4. Quantification of crop sequence variability

Information on land uses in 2013, 2014, 2015 was obtained during the first sampling in 2016 from interviews with landowners and from direct observation of crop residues. Hence, we obtained a five-year crop sequence for each field, which we divided into three three-year sequences (Fig. 1).

**Figure 1.**
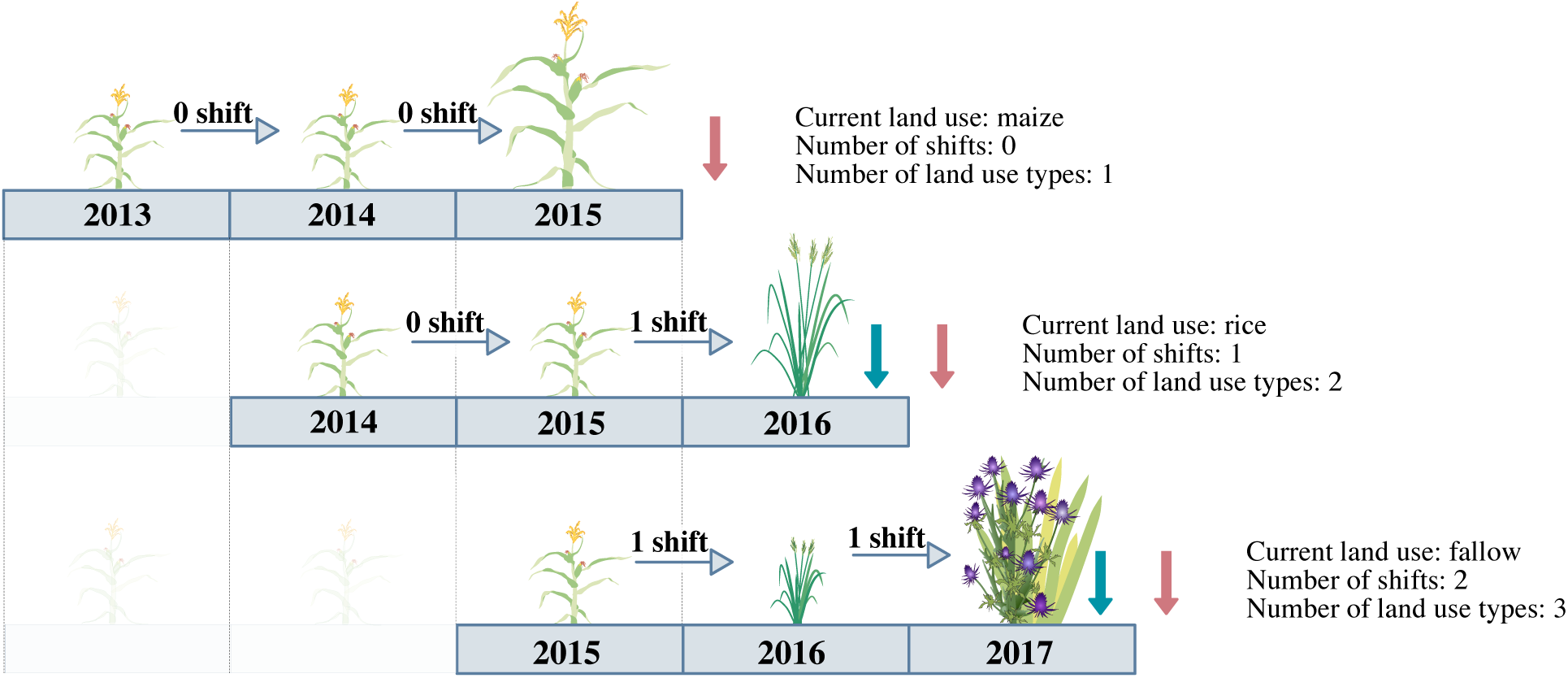
Quantification of the number of crop shifts or the number of crop types, over three growing seasons (example crop sequence: maize - maize - maize - rice - fallow). Larger symbols indicate the crop type at the time of sampling, while smaller symbols indicate previous crops. Vertical arrows indicate the time of sampling: for each cropping season, sampling was conducted in the rainy season just after harvest (blue arrow, same calendar year) and in the dry season, 4-5 months after harvest (red arrow, following calendar year).

We quantified crop sequence in two ways, (1) number of crop shifts over three years, (2) total number of crops involved in the three years. We described the crop using two variables, (1) annual crop type - either maize, upland rice, or no crop i.e. fallow - and (2) young trees intercropping with annual crop or annual crop solo (Table 2). These trees were either rubber or longan trees.

**Table 2.** Description of crop type in the sampled fields (i.e. from March 2016 onwards) based on the variables annual crop and presence of young trees (rubber or longan). In parenthesis: sample size for dry and rainy season.

|  | Maize | Rice | No annual crop |
| --- | --- | --- | --- |
| **No young trees** | Maize monoculture<br>(dry season: n = 12, rainy season n = 8) | Upland rice monoculture (dry season n = 6, rainy season n = 3) | Fallow<br>(rainy season n = 2, dry season n = 2) |
| **Young trees** | Young trees with maize intercrop<br>(dry season n = 6, rainy season n = 1) | Young trees with rice intercrop<br>(dry season n = 8, rainy season n = 8) | Young trees without intercrop<br>(dry season n = 11, rainy season n = 11) |

For instance, maize-maize-maize sequence counts as no shifts and one crop type; maize-maize-rice counts as one shift and two crop types; maize-rice-maize counts as two shifts and two crop types; and maize-rice-fallow counts as two shifts and three crop types. For some fields, we were able to describe crop history only from 2014 onwards, in which case the first sequence was not used (Table 3).

**Table 3.**
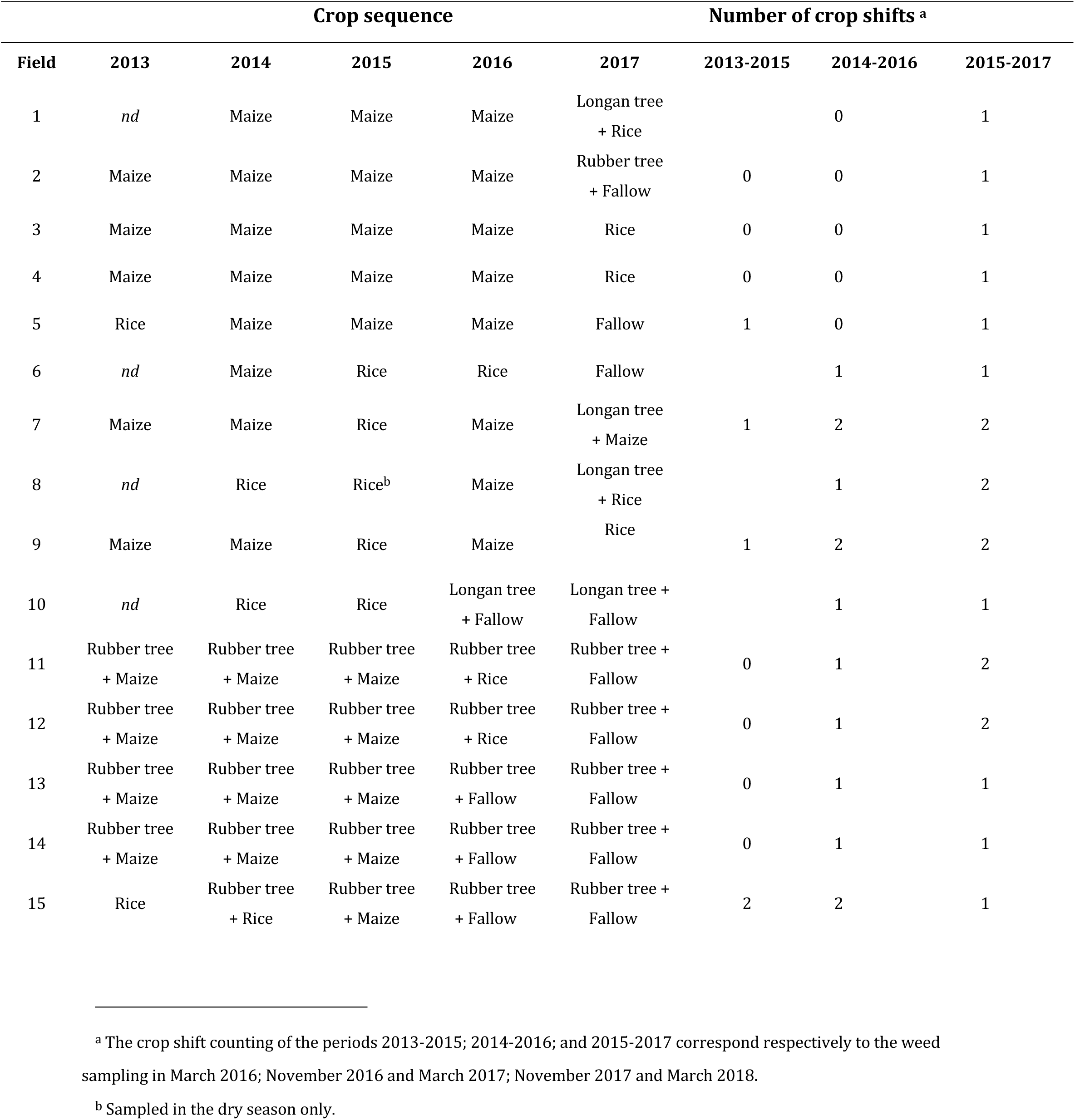
Crop sequences between 2013 and 2017. The number of crop shifts is calculated based on the crop types of the three previous growing seasons (see Fig. 1). *Nd*: no available data: the three-years sequence starting with this year was not included.

| Crop sequence |  |  |  |  |  | Number of crop shifts <sup>a</sup> |  |  |
| --- | --- | --- | --- | --- | --- | --- | --- | --- |
| Field | 2013 | 2014 | 2015 | 2016 | 2017 | 2013-2015 | 2014-2016 | 2015-2017 |
| 1 | *nd* | Maize | Maize | Maize | Longan tree<br>+ Rice |  | 0 | 1 |
| 2 | Maize | Maize | Maize | Maize | Rubber tree<br>+ Fallow | 0 | 0 | 1 |
| 3 | Maize | Maize | Maize | Maize | Rice | 0 | 0 | 1 |
| 4 | Maize | Maize | Maize | Maize | Rice | 0 | 0 | 1 |
| 5 | Rice | Maize | Maize | Maize | Fallow | 1 | 0 | 1 |
| 6 | *nd* | Maize | Rice | Rice | Fallow |  | 1 | 1 |
| 7 | Maize | Maize | Rice | Maize | Longan tree<br>+ Maize | 1 | 2 | 2 |
| 8 | *nd* | Rice | Rice<sup>b</sup> | Maize | Longan tree<br>+ Rice |  | 1 | 2 |
| 9 | Maize | Maize | Rice | Maize | Rice | 1 | 2 | 2 |
| 10 | *nd* | Rice | Rice | Longan tree<br>+ Fallow | Longan tree +<br>Fallow |  | 1 | 1 |
| 11 | Rubber tree<br>+ Maize | Rubber tree<br>+ Maize | Rubber tree<br>+ Maize | Rubber tree<br>+ Rice | Rubber tree +<br>Fallow | 0 | 1 | 2 |
| 12 | Rubber tree<br>+ Maize | Rubber tree<br>+ Maize | Rubber tree<br>+ Maize | Rubber tree<br>+ Rice | Rubber tree +<br>Fallow | 0 | 1 | 2 |
| 13 | Rubber tree<br>+ Maize | Rubber tree<br>+ Maize | Rubber tree<br>+ Maize | Rubber tree<br>+ Fallow | Rubber tree +<br>Fallow | 0 | 1 | 1 |
| 14 | Rubber tree<br>+ Maize | Rubber tree<br>+ Maize | Rubber tree<br>+ Maize | Rubber tree<br>+ Fallow | Rubber tree +<br>Fallow | 0 | 1 | 1 |
| 15 | Rice | Rubber tree<br>+ Rice | Rubber tree<br>+ Maize | Rubber tree<br>+ Fallow | Rubber tree +<br>Fallow | 2 | 2 | 1 |

<sup>a</sup> The crop shift counting of the periods 2013-2015; 2014-2016; and 2015-2017 correspond respectively to the weed sampling in March 2016; November 2016 and March 2017; November 2017 and March 2018.
<sup>b</sup> Sampled in the dry season only.

Maize was the most represented annual crop in the dataset. In order to validate our main results regarding the impact of crop shifts irrespective of the identity of the current crop, all analyses were conducted twice: i/ including all available data; and ii/ including only fields with maize as the current annual crop (i.e. maize monoculture or young tree plantations with maize intercrop). For these analyses, we compared maize fields with no crop shift (n = 17) or two crop shifts (n = 7) because there were not enough maize fields with only one shift (n = 3).

### 2.5. Statistical analyses

All analyses were conducted using R (R Core Team, 2018). We built models for each weed community characteristic described in Table 1 as a response variable. We used linear mixed models (functions *lmer* and *lme*, packages lme4 (Bates et al., 2014) and nlme (Pinheiro et al., 2018) in R). Each model included as explanatory variables: the annual crop type, the presence of trees, and a spatial covariate (see below), and the temporal crop variability and its interaction with sampling season. The temporal crop variability was measured either as the number of crop shifts (levels: 0, 1 or 2) or the total number of crops involved (1, 2 or 3) in the three preceding sampling seasons (2 df, Degrees of freedom).

Spatial autocorrelation was also included in the model, fitted with the *lme* function, as an exponential correlation structure (function *corSpatial).* Beforehand we scaled the easting and northing coordinates of each field and added a small jitter (normal noise, mean of 0 and standard deviation of 0.001) because the function cannot handle duplicate coordinates. Other forms of spatial autocorrelation (gaussian, linear, spherical) were tested but model AIC were similar. All significance values and estimates are extracted from these models. However, to provide estimates of partial R^2^ for each single variable (function *r2beta*, package r2glmm, Kenward Roger method, Jaeger (2017)) we had to use model fitted with the *lmer* function (i.e. without spatial autocorrelation) due to package compatibility issues. As the autocovariates were not significant, this was unlikely to strongly affect the R^2^.

Besides, the treatment of interest (number of crop shifts) was not independent of the field itself or the crop type. Indeed, we conducted botanical inventories repeatedly in the same fields and the number of crop shifts in one site was not independent from one year to another: a field with two crop shifts (or respectively zero) on a given year would necessarily have at least (resp. at most) one shift the next year. In order to take into account this non-independence, we used field-level random effects in the models. Plant densities (always strictly positive) were log-transformed, and biomasses square-root-transformed, to ensure normality of the residuals. We conducted pairwise comparisons between each level of crop temporal variability (e.g., for the number of crop shifts, 0 v. 1, 0 v. 2 and 1 v. 2 shifts; and for the number of crop types, 1 v. 2, 1 v. 3, and 2 v. 3 crop types) while keeping the other explanatory variables constant (*emmeans* function, package emmeans, Lenth (2018)). The significance of each variable in the full model was assessed using Anova type II or III tests (function *Anova*, package car, Fox et al. (2011)): thus, the effect of each variable was tested “after”, or “controlling for” the other fixed effects present in the model.

Models including only maize fields were also mixed models including spatial autocovariates, and the explanatory variables were the same (except for the annual crop variable, which was not included).

In additional sensitivity analyses, the year was also included as a fixed effect in the models. The models were then selected based on AIC. The results from these analyses were consistent with our main results and are not presented for concision.

### 2.6. Indicator species analysis

Our last objective was to determine whether certain species were specifically associated with either high or low frequency of crop shifts. In this regard, we identified groups of indicator species related to each number of crop shifts. As proposed by Cáceres and Legendre (2009), indicator species are species that can be used as ecological indicators of environmental and ecological conditions. Their association to a given environment (here, over periods of three years, the number of crops or number of shifts) is based on the specificity and the fidelity of the species as an indicator of the environmental group (i.e. here, fields with a given frequency of crop shifts). We used the indicspecies package (function *multipatt*, IndVal.g method, Cáceres and Legendre (2009)).

## 3. Results

### 3.1. Diversification of crop sequences

From 2014 to 2018, we observed a total of 22 distinct crop sequences (Fig. 2), and an increasing diversity in crop sequences with time. In the period 2013-2015 five different sequences occurred, in the period 2015-2017 this increased to eleven (Fig 2). For instance, continuous maize cultivation (i.e. the sequence maize - maize – maize) was only observed in the period 2015-2017. Instead, fallows appeared and young rubber tree plantations were intercropped with rice or maize during four to five years before the shade from the canopy prevented further intercropping.

**Figure 2.**
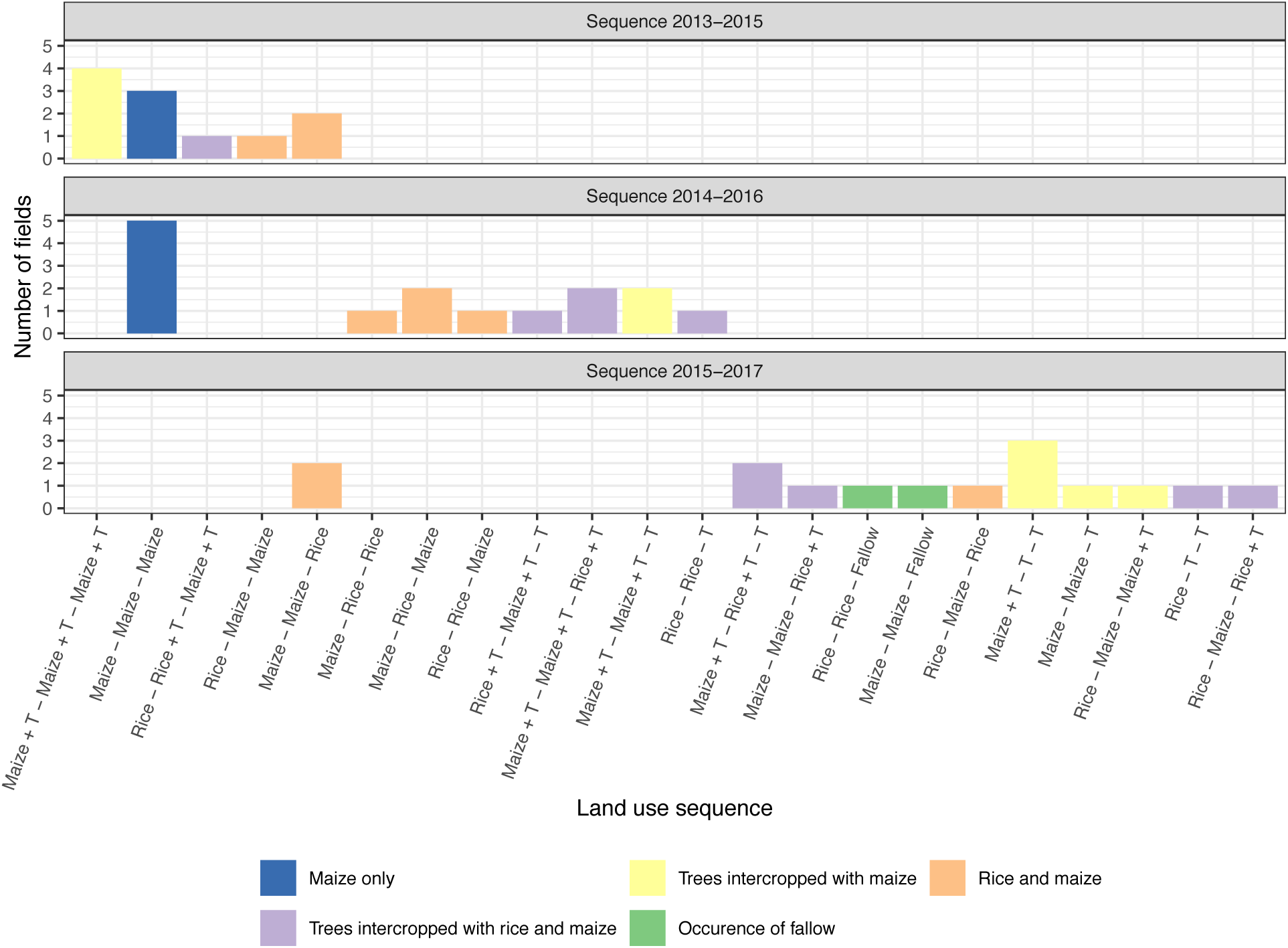
Frequency distribution of 22 combinations of crop sequences occurring in farmers’ fields over the period 2013-2017. T indicates the presence of young trees, i.e. rubber or longan.

The number of crop shifts was not independent of crop type: maize fields were more often associated with continuous cropping (no shift) compared to upland rice or fallows which were always subject to crop shifts after 1 or 2 years (**X**^2^ test, P < 10^6^). Some farmers reported that the decision to grow maize (cash crop) or rice (subsistence crop) depended on both market and familial factors: a family running out of rice would be more likely to grow rice the next year, while the market price of maize might determine the planting of maize fields. This decision was also likely to depend on the start of the rainy season, as maize has a shorter cycle and needs therefore only part of the rainy season to complete its growth.

### 3.2. Plant richness

We found a total of 56 herbaceous species (52 identified at least to genus level), and 79 woody species (all identified) (Table S1). Fields in the dry and rainy season were equally species-rich, with 6 to 26 (median 18) species per field in the dry season and 7 to 32 (median 18) species per field in the rainy season (P = 0.2, after controlling for crop type and crop shifts). We identified three groups of species based on breaks in the Relative Importance index bar plot (Fig. 3). Three herbaceous weeds (*Ageratum conyzoides* and *Conyza sumatrensis*, Asteraceae and *Mitracarpus hirtus*, Rubiaceae) had a Relative Importance Index > 0.4, indicating that they were both very frequent and abundant. We identified 9 intermediate species with a RI comprised between 0.2 and 0.4. The least common species had a RI lower than 0.1 and comprised most shrub and tree species; among them, 43 species had a RI < 0.01.

**Figure 3.**
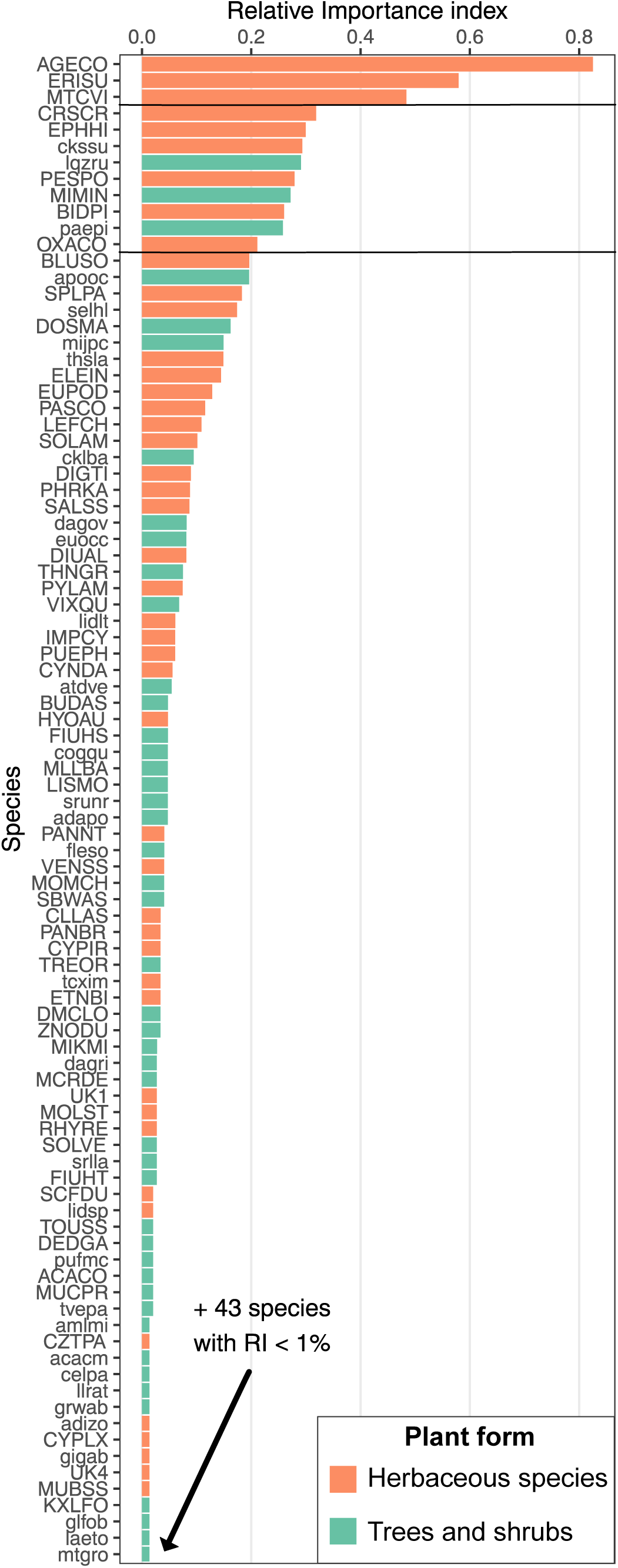
Relative Importance index divides the identified species into three groups: (1) dominant, ubiquitous species, here *A. conyzoides* (AGECO), *C. sumatrensis* (ERISU) and *M. hirtus* (MTCVI) (top part) (2) intermediate species (middle part) and (3) rare species (bottom part). Species codes can be found in Table S1.

Figure 4 shows an increase in species number among herbaceous species with more frequent crop shifts, particularly in the dry season. In fields where the crop changed every year, an average of 13 herbaceous species were recorded in the third year. This is a 36 % increase (P = 0.03), compared to fields where three years the same crop was cultivated, where only 8.3 herbaceous species were recorded on the third year (Fig. 4). This effect was stronger than that of the crop type itself (based on partial R^2^, Table 4). The richness of trees and shrubs species or of the overall community was not affected by the number of crop shifts either during the dry or the rainy season (Fig. 4, Table 4). When considering the number of crop types instead of the number of crop shifts, herbaceous species number also tended to increase with the number of crop types but the relationship was not significant.

**Table 4.**
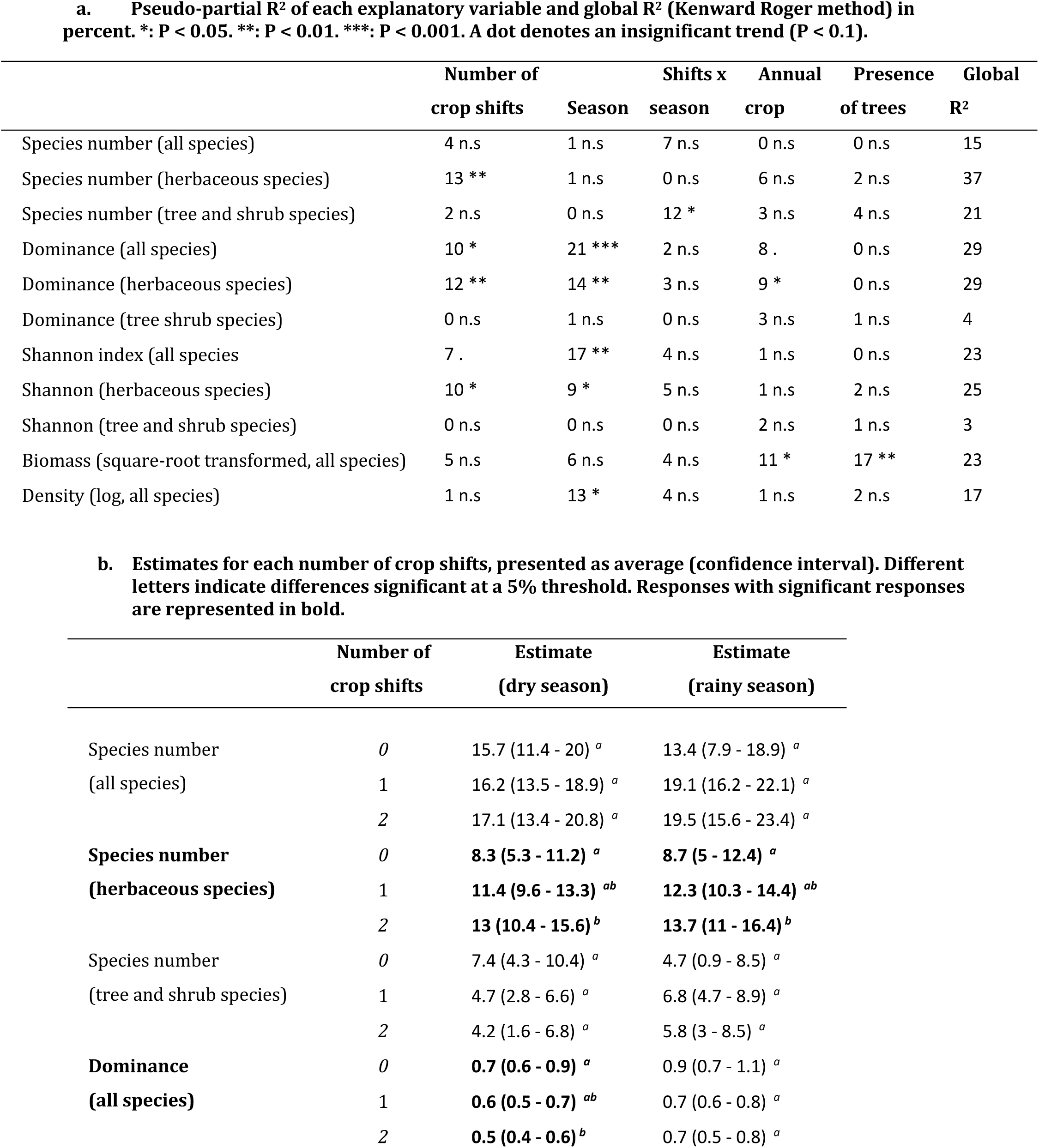
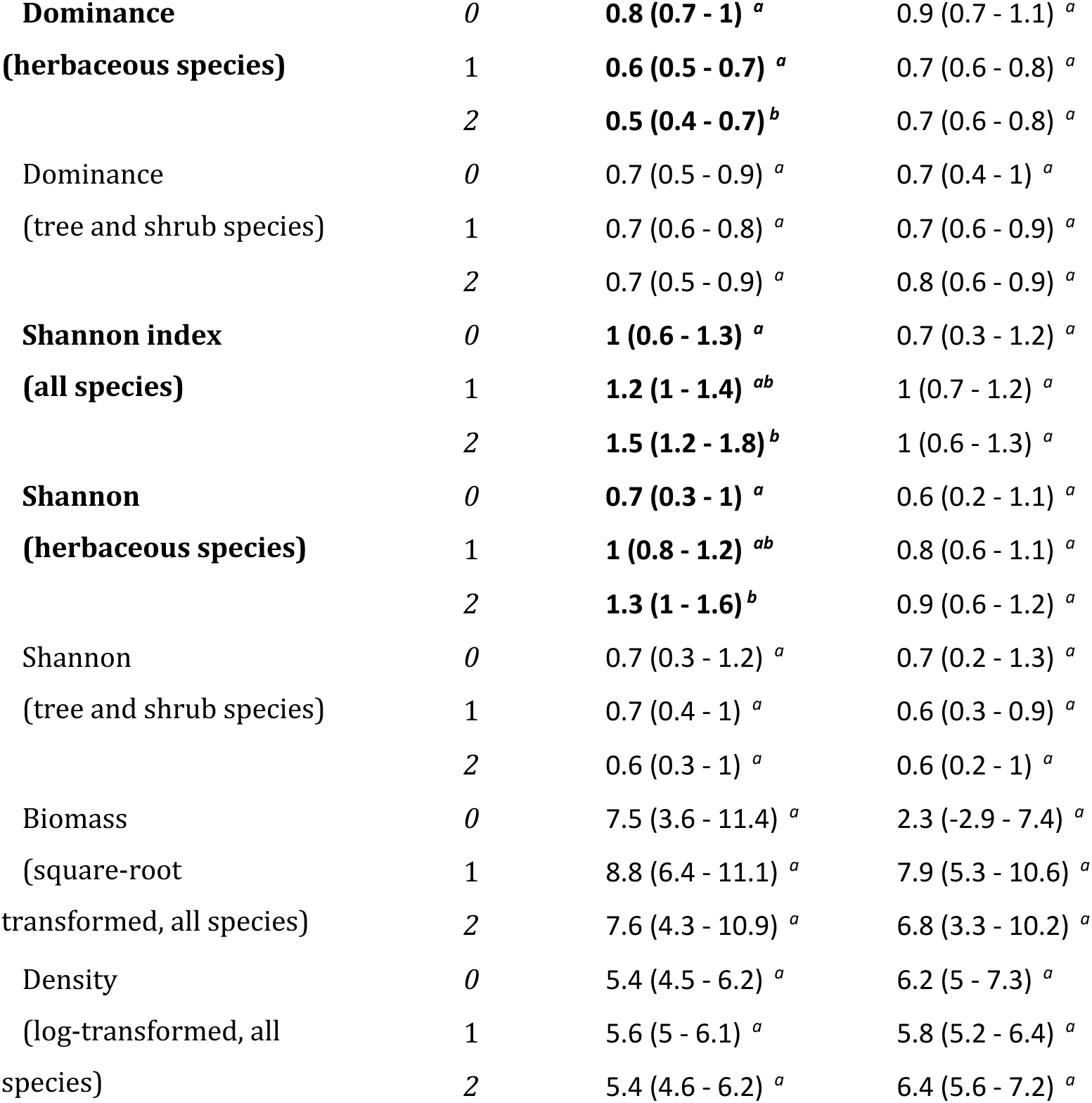
Effects of crop type and crop shifts on weed richness, diversity and abundance: results from mixed model analyses.

|  | Number of<br>crop shifts | Season | Shifts x<br>season | Annual<br>crop | Presence<br>of trees | Global<br>R<sup>2</sup> |
| --- | --- | --- | --- | --- | --- | --- |
| Species number (all species) | 4 n.s | 1 n.s | 7 n.s | 0 n.s | 0 n.s | 15 |
| Species number (herbaceous species) | 13 ** | 1 n.s | 0 n.s | 6 n.s | 2 n.s | 37 |
| Species number (tree and shrub species) | 2 n.s | 0 n.s | 12 * | 3 n.s | 4 n.s | 21 |
| Dominance (all species) | 10 * | 21 *** | 2 n.s | 8 . | 0 n.s | 29 |
| Dominance (herbaceous species) | 12 ** | 14 ** | 3 n.s | 9 * | 0 n.s | 29 |
| Dominance (tree shrub species) | 0 n.s | 1 n.s | 0 n.s | 3 n.s | 1 n.s | 4 |
| Shannon index (all species) | 7 . | 17 ** | 4 n.s | 1 n.s | 0 n.s | 23 |
| Shannon (herbaceous species) | 10 * | 9 * | 5 n.s | 1 n.s | 2 n.s | 25 |
| Shannon (tree and shrub species) | 0 n.s | 0 n.s | 0 n.s | 2 n.s | 1 n.s | 3 |
| Biomass (square-root transformed, all species) | 5 n.s | 6 n.s | 4 n.s | 11 * | 17 ** | 23 |
| Density (log, all species) | 1 n.s | 13 * | 4 n.s | 1 n.s | 2 n.s | 17 |

|  | Number of<br>crop shifts | Estimate<br>(dry season) | Estimate<br>(rainy season) |
| --- | --- | --- | --- |
| Species number | 0 | 15.7 (11.4 - 20) <sup>a</sup> | 13.4 (7.9 - 18.9) <sup>a</sup> |
| (all species) | 1 | 16.2 (13.5 - 18.9) <sup>a</sup> | 19.1 (16.2 - 22.1) <sup>a</sup> |
|  | 2 | 17.1 (13.4 - 20.8) <sup>a</sup> | 19.5 (15.6 - 23.4) <sup>a</sup> |
| **Species number** | 0 | **8.3 (5.3 - 11.2) <sup>a</sup>** | **8.7 (5 - 12.4) <sup>a</sup>** |
| **(herbaceous species)** | 1 | **11.4 (9.6 - 13.3) <sup>ab</sup>** | **12.3 (10.3 - 14.4) <sup>ab</sup>** |
|  | 2 | **13 (10.4 - 15.6) <sup>b</sup>** | **13.7 (11 - 16.4) <sup>b</sup>** |
| Species number | 0 | 7.4 (4.3 - 10.4) <sup>a</sup> | 4.7 (0.9 - 8.5) <sup>a</sup> |
| (tree and shrub species) | 1 | 4.7 (2.8 - 6.6) <sup>a</sup> | 6.8 (4.7 - 8.9) <sup>a</sup> |
|  | 2 | 4.2 (1.6 - 6.8) <sup>a</sup> | 5.8 (3 - 8.5) <sup>a</sup> |
| **Dominance** | 0 | **0.7 (0.6 - 0.9) <sup>a</sup>** | 0.9 (0.7 - 1.1) <sup>a</sup> |
| **(all species)** | 1 | **0.6 (0.5 - 0.7) <sup>ab</sup>** | 0.7 (0.6 - 0.8) <sup>a</sup> |
|  | 2 | **0.5 (0.4 - 0.6) <sup>b</sup>** | 0.7 (0.5 - 0.8) <sup>a</sup> |
| **Dominance** | 0 | **0.8 (0.7 - 1) <sup>a</sup>** | 0.9 (0.7 - 1.1) <sup>a</sup> |
| **(herbaceous species)** | 1 | **0.6 (0.5 - 0.7) <sup>a</sup>** | 0.7 (0.6 - 0.8) <sup>a</sup> |
|  | 2 | **0.5 (0.4 - 0.7) <sup>b</sup>** | 0.7 (0.6 - 0.8) <sup>a</sup> |
| Dominance | 0 | 0.7 (0.5 - 0.9) <sup>a</sup> | 0.7 (0.4 - 1) <sup>a</sup> |
| (tree and shrub species) | 1 | 0.7 (0.6 - 0.8) <sup>a</sup> | 0.7 (0.6 - 0.9) <sup>a</sup> |
|  | 2 | 0.7 (0.5 - 0.9) <sup>a</sup> | 0.8 (0.6 - 0.9) <sup>a</sup> |
| **Shannon index** | 0 | **1 (0.6 - 1.3) <sup>a</sup>** | 0.7 (0.3 - 1.2) <sup>a</sup> |
| **(all species)** | 1 | **1.2 (1 - 1.4) <sup>ab</sup>** | 1 (0.7 - 1.2) <sup>a</sup> |
|  | 2 | **1.5 (1.2 - 1.8) <sup>b</sup>** | 1 (0.6 - 1.3) <sup>a</sup> |
| **Shannon** | 0 | **0.7 (0.3 - 1) <sup>a</sup>** | 0.6 (0.2 - 1.1) <sup>a</sup> |
| **(herbaceous species)** | 1 | **1 (0.8 - 1.2) <sup>ab</sup>** | 0.8 (0.6 - 1.1) <sup>a</sup> |
|  | 2 | **1.3 (1 - 1.6) <sup>b</sup>** | 0.9 (0.6 - 1.2) <sup>a</sup> |
| Shannon | 0 | 0.7 (0.3 - 1.2) <sup>a</sup> | 0.7 (0.2 - 1.3) <sup>a</sup> |
| (tree and shrub species) | 1 | 0.7 (0.4 - 1) <sup>a</sup> | 0.6 (0.3 - 0.9) <sup>a</sup> |
|  | 2 | 0.6 (0.3 - 1) <sup>a</sup> | 0.6 (0.2 - 1) <sup>a</sup> |
| Biomass | 0 | 7.5 (3.6 - 11.4) <sup>a</sup> | 2.3 (-2.9 - 7.4) <sup>a</sup> |
| (square-root | 1 | 8.8 (6.4 - 11.1) <sup>a</sup> | 7.9 (5.3 - 10.6) <sup>a</sup> |
| transformed, all species) | 2 | 7.6 (4.3 - 10.9) <sup>a</sup> | 6.8 (3.3 - 10.2) <sup>a</sup> |
| Density | 0 | 5.4 (4.5 - 6.2) <sup>a</sup> | 6.2 (5 - 7.3) <sup>a</sup> |
| (log-transformed, all | 1 | 5.6 (5 - 6.1) <sup>a</sup> | 5.8 (5.2 - 6.4) <sup>a</sup> |
| species) | 2 | 5.4 (4.6 - 6.2) <sup>a</sup> | 6.4 (5.6 - 7.2) <sup>a</sup> |

**Figure 4.**
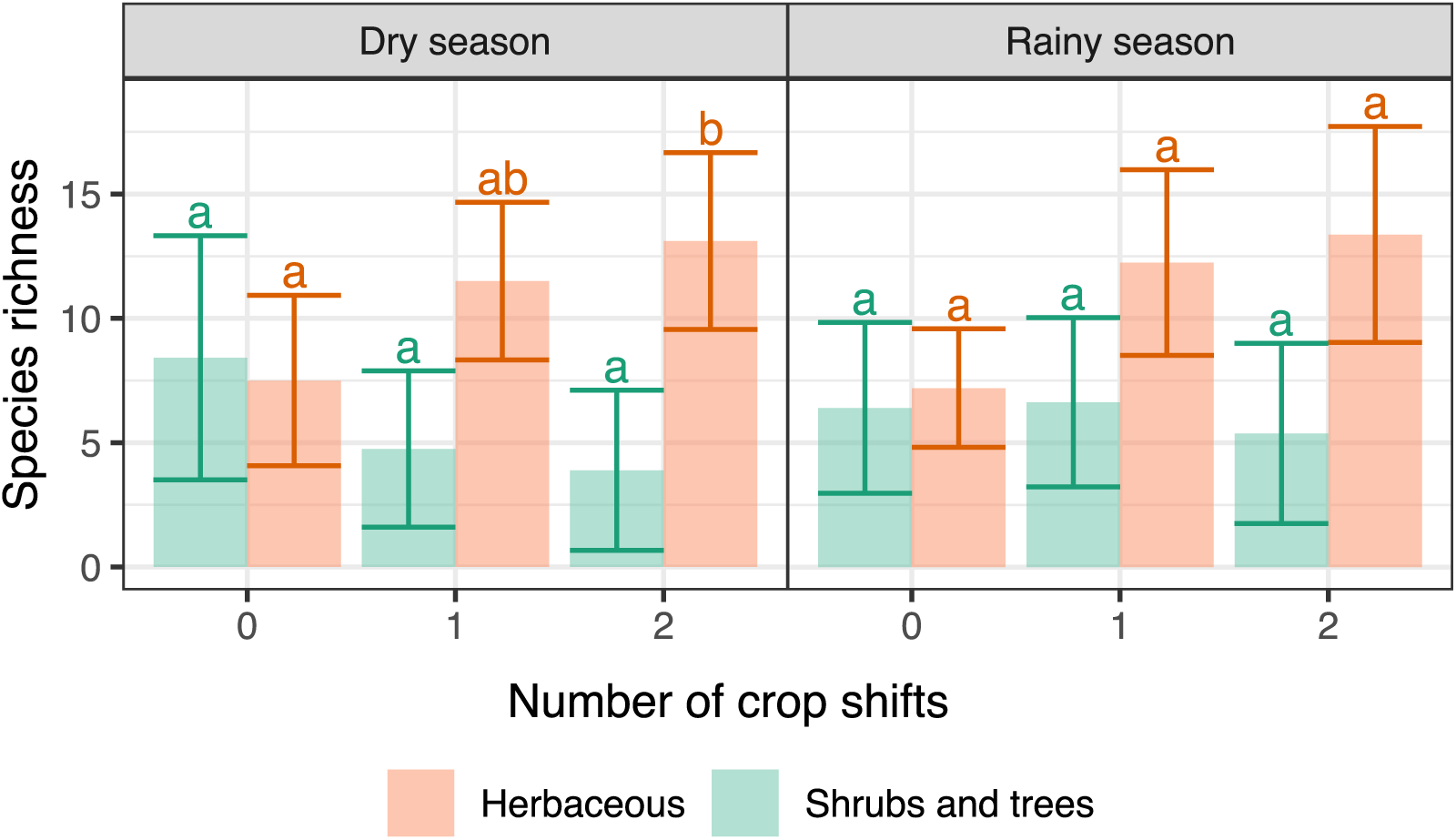
Relationships between species richness per field and number of crop shifts. Bars represent the mean + /- standard deviation, different letters indicate significant differences (P < 0.05) within each group (herbaceous or shrub/trees). a) Dry season samples, b) Rainy season samples.

Similarly, maize monocultures tended to have lower herbaceous species number than fields with maize as the current annual crop (i.e. maize fields or young tree plantations intercropped with maize) with two shifts in the three previous years. It was significant in the rainy season (P = 0.02) but only a trend in the dry season (P = 0.07, Table S3, Fig. S1).

### 3.3. Plant biomass and density

Plant biomass (square root-transformed) and density (log-transformed) did not vary with the number of crop shifts nor the number of crop types (P > 0.1, Table 4, S2). When controlling for the number of shifts, annual crop type and the presence of trees, weed communities had lower densities (228 plant/m^2^) and slightly higher biomass (70.2 g/m^2^) in the dry compared to the rainy season (density of 412 plant/m^2^ (P = 0.01) and biomass of 44 g/m^2^ (N.S. P = 0.08)).

Maize monocultures had significantly lower plant biomass (3.2 g/m^2^ on average) and plant density (403 plant/m^2^) than maize fields with two shifts in the three previous years (respectively 88 g/m^2^, P = 0.01; and 1636 plant/m^2^; P = 0.03) in the rainy season. The difference was not significant in the dry season (P > 0.3, Table S3).

### 3.4. Plant diversity

Herbaceous weed communities were more diverse when the number of crop shifts increased, particularly in the dry season. There, the Shannon index H’ increased by 46% from fields with no crop shift (0.7 on average) to in fields with two crop shifts (1.3, P = 0.009, Table 4). This effect was stronger than that of the crop type itself (based on partial R^2^, Table 4). The same trend was observed for diversity in the total community (shrubs/trees included, P = 0.04) but not for shrub/tree communities only (P > 0.8). Herbaceous plant communities were generally more diverse in the dry season (H’ = 0.98) compared to rainy season (H’ = 0.78, P = 0.03).

All communities were dominated by the two most abundant species – usually *A. conyzoides* (dominant in 57% of the fields) followed by *C. sumatrensis* (22% of the fields) -, yet this dominance was weaker when the number of crop shifts increased. Indeed, in fields with no shift, the dominance (Berger-Parker index, D) was 0.8 on average, which indicates that the most abundant species represented 80 % of all individuals. Conversely, dominance index decreased by 38% (to D = 0.5 on average) in fields with two crop shifts (P = 0.03, Table 4). Similar trends for diversity and dominance were observed when considering the number of crops (Table S2).

When controlling for the crop shifts, crop type and the presence of trees, dominance of herbaceous species, but not of shrubs and trees was significantly higher in the rainy (D = 0.7) than the dry season (D = 0.6, P < 0.05, data not shown). This can be explained by the declining numbers of the two dominant weeds *A. conyzoides* and *Conyza sumatrensis*, both annual species. Annual species tend to end their life cycle in the dry season, modifying dominance and diversity. This was not observed in woody species because they are all perennials. Conversely, Shannon index was higher in the dry (H’ = 0.98) compared to rainy season (H’ = 0.78, P = 0.03) for herbaceous species; it was also significant for all species but not for shrub and tree species (data not shown).

Consistently with the results considering all crop types, the diversity of the total community in the dry season was lower in maize monocultures (H’ = 1.1) than in maize fields with two shifts (H’ = 1.6, P = 0.01, Table S3). Diversity for the total community followed the same trend as for herbaceous species only, probably because the woody fraction was altogether relatively small in maize monoculture. Dominance did not vary significantly between maize monocultures and maize with two shifts.

### 3.5. Indicator species

We found that 12 species were indicators of a precise number of crop shifts, or a range of crop shifts, in the three years preceding sampling (Table 5). For example, *Digitaria radicosa* (annual, creeping C4 grass) and *Centella asiatica* (perennial, creeping herb) were strongly associated to fields with two shifts while the shade-tolerant fern *Thelopterys subelatus* was a good indicator of fields with either one or two crop shifts in the previous three years. Conversely, *Antidesma velutinosum*, a tree, *Streblus asper*, a shrub and *Tournefortia sp.*, a liana, were found mostly in continuous cropping, with no shifts over the previous three years.

**Table 5.** Indicator species associated with fields with zero, one or two crop shifts in the three years preceding sampling. Test statistic is the IndVal.g association index. *** P < 0.001, ** P < 0.01, * P < 0.05.

| Growth form | Species | Test statistic | Significance |
| --- | --- | --- | --- |
| **Fields with zero shift** |  |  |  |
| Tree | *Antidesma velutinosum* | 0.534 | * |
| Shrub | *Streblus asper* | 0.485 | * |
| Liana | *Tournefortia sp.* | 0.420 | * |
| **Fields with zero to one shift** |  |  |  |
| Tree | *Diospyros malabarica* | 0.629 | * |
| Large liana | *Millettia pachycarpa* | 0.599 | * |
| **Fields with one to two shifts** |  |  |  |
| Ground fern | *Thelopterys subelatus* | 0.830 | *** |
| Annual herb | *Bidens pilosa* | 0.749 | * |
| Perennial C<sub>3</sub> grass | *Thysanolaena latifolia* | 0.625 | * |
| **Fields with two shifts** |  |  |  |
| Annual C<sub>4</sub> grass | *Digitaria radicata* | 0.547 | ** |
| Tree | *Vitex quinata* | 0.510 | * |
| Perennial herb | *Centella asiatica* | 0.479 | ** |
| Perennial vine | *Thumbergia grandiflora* | 0.475 | * |

## 4. Discussion

### 4.1. Crop sequence variability

In northern Thailand farmers often grow maize for several consecutive years, with occasionally upland rice to break the maize monoculture. Conversely, upland rice is only cultivated for 1 or 2 consecutive years. Intercropping of upland rice and maize with young rubber trees whose canopy is still open enough or with longan saplings is another option for farmers. The famers’ motives for crop shifts and intercropping were difficult to apprehend and impossible to verify. However, we can distinguish between socio-economic reasons and ecological-technical ones.

Socio-economic motives include market prices and labour availability. For instance, the persistently low market price for rubber since 2011 induces farmers to plant fruit trees (longan) instead of rubber trees. Labour demand in upland rice is more important than in maize, yet rice requires less modern inputs compared to maize, e.g. hybrid seed, frequent herbicide spraying. More interesting, because linked to weed control strategies, are the ecological and technical motives for crop sequences. Weed infestation usually prevents cultivating rice more than 2-3 years in a row while maize as a crop is much more resistant to weeds (Sankaran and de Datta, 1986). Moreover, the herbicide-resistant maize varieties allow the continuing use of herbicides later in the season. In the study area weeds do not seem to limit the continuous cultivation of maize, however, a growing built-up of aggressive weeds could compromise a shift to upland rice. For instance, heavily infested fields in Laos no longer support upland rice (Dupin *et al*, 2009). The local bimodal rainfall distribution has a long rainy season but is subject to intermittent dry spells, which could lead farmers to favour maize over upland rice. Indeed, sowing and harvest dates of the traditional long-cycle upland rice varieties cannot be modified, while the short-cycle hybrid maize is more flexible: for instance, we observed both harvested and immature maize fields during our October 2016 sampling. The reason for early or late planting could be to avoid drought. A second advantage of maize is that the hill density can be reduced to adjust to water availability: low density planting provides more opportunities for weeds to grow, but weeds can be controlled by late herbicide spraying. In upland rice, hill density is high and cannot be modified without compromising the weed-competitiveness of the leafy, high stature varieties.

This difference between rice and maize cultivation led to the emergence of two groups of crop sequences among the fields we investigated. One consisted of continuous maize cultivation (either continuous maize monoculture or intercrop between tree rows) and the other with alternating cultivation of maize and rice. Neyret *et al.* (2018) reported higher richness and greater diversity of weed communities in upland rice than in maize. A similar result is demonstrated here, in crop sequences containing upland rice and in sequences of only maize. Differences in cultivation and weeding practices among crops are at least partly responsible for the crop-effect in weed communities (Neyret et al., 2018). This leads to variability in community composition from year to year and thus to the increase of weed richness with the number of crop shifts. Thus, while the crop type effect and the “number of crop shifts” effect are partly confounded in this study, we tried to separate the two effects, i.e. by (1) including the effect of annual crop type and the presence of trees in the models, and (2) controlling for crop type before measuring the effect of the number of shifts. Besides, the positive effect of crop shifts on weed richness was at least partly supported when considering only maize fields, i.e. when removing any possible confusion with the annual crop currently grown in the field. Thus, while crop type was very likely to affect the richness, diversity and abundance of weeds in this trial, we showed that the number of shifts had an effect independently from crop type.

### 4.2. Crop shifts increase herbaceous species number and diversity

Our finding that plant diversity increases with shift frequency is also consistent with the literature. Previous studies, mostly in temperate areas, indeed found that increasing the inter-annual variability of land use had a positive effect on weed richness and diversity (Liebman and Dyck, 1993; Doucet et al., 1999; Squire et al., 2000; Ulber et al., 2009). For instance, Allan et al. (2014) showed that in grasslands, weed richness (and especially rare plants richness) increased with the temporal heterogeneity of fertilisation, mowing and grazing intensity, independently of the level of intensity itself. Using a simulation approach, Bürger et al. (2015) showed that while tillage was the main factor affecting weed diversity, the simplification of crop rotations also reduced biodiversity, especially in regions already harbouring low diversities.

In addition, our study showed that the effect of the number of crop shifts on herbaceous species number and diversity was stronger than that of crop type, contrarily to previous results (Bàrberi et al., 1997; Smith and Gross, 2007). The outcome of studies investigating the effects of temporal diversity on plants depends on the timescale of the study. Such effects are likely to be noticeable only when looking at the total weed flora within a field, by looking either at the seedbank or at the flora over multiple years, as opposed to looking at the flora within a single year only (Dessaint et al., 1997). Altogether, this suggests that in the long-term, the severely degrading effect on biodiversity of some crops could be offset by annual rotations with more biodiversity-friendly land uses. Additional studies of similar agro-ecosystems investigating the effect of longer-term crop rotations could thus provide further confirmation of our results.

### 4.3. Species associated with high shift frequency

Although no species were entirely restricted to a given frequency of crop shifts, we were able to show that some species were significantly associated to either frequently shifting fields or continuous crops. The three dominant herbaceous species – *Ageratum conyzoides, Mitracarpus hirtus* and *Conyza sumatrensis*) were ubiquitous, and thus not associated to any particular crop. All species associated with fields with zero shift or one to two shifts were tree species, while the two species most significantly associated with fields with two shifts (*Digitaria radicosa* and *Centella asiatica*) were herbaceous annuals. Besides, contrarily to herbaceous species, trees and shrubs richness and diversity did not respond to changes in the number of crop shifts or the number of land use types. This suggests a weaker response of trees and shrubs to year-to-year shifts, compared to herbaceous species, which grow and reproduce more quickly. Indeed, perennial species (including a few of the herbaceous species, but all shrub and tree species) have more underground reserves from which they can directly regrow (Raunkiaer, 1934). This makes them less dependent on local conditions and farming practices to establish in a given field.

### 4.4. Seasonal effect on weeds richness and abundance

We showed that although species number did not vary significantly with season, there was a strong decrease in plant biomass, but increase in plant density in the rainy season compared to the dry season. Fewer yet larger individual plants in the dry season than in the rainy season can be explained by a combination of at least three factors. The life cycle of annual weeds runs with the rainy season so their life cycle naturally ends in the dry season, reducing density. Secondly, the dry season, for its lack of surface soil moisture, is less suitable than the rainy season for new emergences; instead, well-established individuals can expand in the dry season, their roots exploring moisture in the deeper soil layers. Finally, the fields sampled in the dry season had not experienced weed control for at least six months, allowing for self-thinning among seedlings and the outgrow of the more vigorous individuals. Most of the seedlings found in the rainy season belonged to the three dominant species, which also explains the decrease in species diversity, and the increase in the dominance index. *A. conyzoides* in particular has a very effective reproduction rate, producing numerous seeds with high germination rates (Kohli et al., 2006; Hao et al., 2009). This dominance by the main species in the rainy season was likely to explain the strongest response to crop shifts on species number in the dry compared to rainy season.

### 4.5. Resource and disturbance variability

Gaba et al. (2014) identified two main gradients driving the abundance and diversity of weeds in crop rotations. On the one hand, the resource variability gradient represents the temporal variability of resource availability in the field, which is expected to increase weed diversity through niche diversification. On the other hand, the disturbance gradient represents the type and frequency of disturbance, which is expected to increase mortality rates and to decrease weed abundance.

Contrarily to our expectations, we did not detect a change in weed biomass or weed density with crop shifts. This was probably due to the fact that fields with frequent shifts did not necessarily have a higher variability of disturbance types and timing, which is expected to be the main driver of the changes in weed abundance (Gaba et al., 2014). Maize and rice fields had relatively similar soil preparation as well as fertilisation rates: thus, the type and timing of disturbance were unlikely to differ among fields (except for fallows). While in this trial we did not have data on crop yield, the experiment was farmer-managed, thus weed control was assumed sufficient in achieving an acceptable yield. Additional measures of yield and crop biomass would be needed to confirm whether the frequency of crop shifts affects the actual aggressiveness of weed communities towards the crop. In terms of resource variability, rice and maize created different light conditions, which is known to be an important determinant of weed growth (Holt, 1995): while rice grows very densely, quickly covering the ground and limiting weed growth, maize (planted at relatively low density in the study system) leaves most of the soil bare and triggers the germination of photosensitive species. Many common tropical weed seeds require full sunlight to germinate (example *A. conyzoides* and *C. sumatrensis*), while other prefer light shade (*Chromolaena odorata*) (Garwood, 1989; de Rouw et al., 2013). Thus, in fields with frequent shifts, the reproduction rates of very competitive, heliophilous species with high seed production are regularly lowered by less favourable conditions. This creates opportunities for new species to germinate from the seedbank or to establish from neighbour communities, and explains the lower dominance index in fields with frequent shifts. Frequent shifts thus prevent the selection of species functionally close to the crop (Liebman and Dyck, 1993; Smith et al., 2010).

Sites with permanent annual cropping have large seed banks, between 5000 and 10 000 viable seeds/m^2^ (Garwood, 1989). In Laos, soil seed banks were found similar across upland rice sites, in density and composition, but weed species abundances in the cultivated fields were not correlated with densities in the seed bank. These results indicate that emergence during the cultivation period reflected the local growing conditions far more than their availability in the seed bank (de Rouw et al., 2013). In our study system, rice and maize residues are likely to create different humidity conditions, which could favour the germination of different fractions of the seedbank. Rice and maize also had different sowing and harvesting times, which have been shown to be major determinants of the functional composition of weed communities (Gunton et al., 2011). For instance, maize has a much shorter growing period, which leaves the fields almost fallow-like with dry maize stalks during a large part of the year. The later application of herbicides in maize (e.g. after germination), repeated in maize fields with no shifts, might also have led to the selection of species able to recover quickly after spraying, such as *A. conyzoides*. Thus, crops changing from year to year provide variable germinating and growing conditions for weeds, and a selection of different species from one season to another. This allows the maintenance of diverse communities over time by allowing species with different responses to the environment to coexist stably in different niches (Allan et al., 2014; Gaba et al., 2014).

## 6. Conclusion

Promoting diversified and less competitive weed communities favours the continued provision of weed ecosystem services, such as support for diversity at higher trophic levels or erosion mitigation. By measuring the short-term frequency of crop shifts, we were able to show that the number of crop shifts had a significantly positive effect on the richness and diversity of herbaceous weed communities. It did not, however, affect their overall abundance. These results show that yearly crop shifts in this area are not adequate to control weed biomass and their competitivity toward the crop, but support plant diversity conservation. Future research should address crop shifts in longer and diversified crop sequences in these threatened, rapidly changing agro-ecosystems to further determine their potential for weed control and diversity conservation.

## Acknowledgements

This study was realised during a research project of Sorbonne University and Institut de Recherche pour le Développement; and was supported by the ANR HeveAdapt project (grant ANR-14-CE030012-04) of the French Agence Nationale de la Recherche. Fieldwork was realised with the cooperation of the Huai Lang Royal Project Center and the Land Development Department of Thailand. We thank two anonymous reviewers for their thoughtful comments.

## Data availability

Data and R code used in this study will be made available online upon acceptance.

## Supplementary Material

**Table S1.**
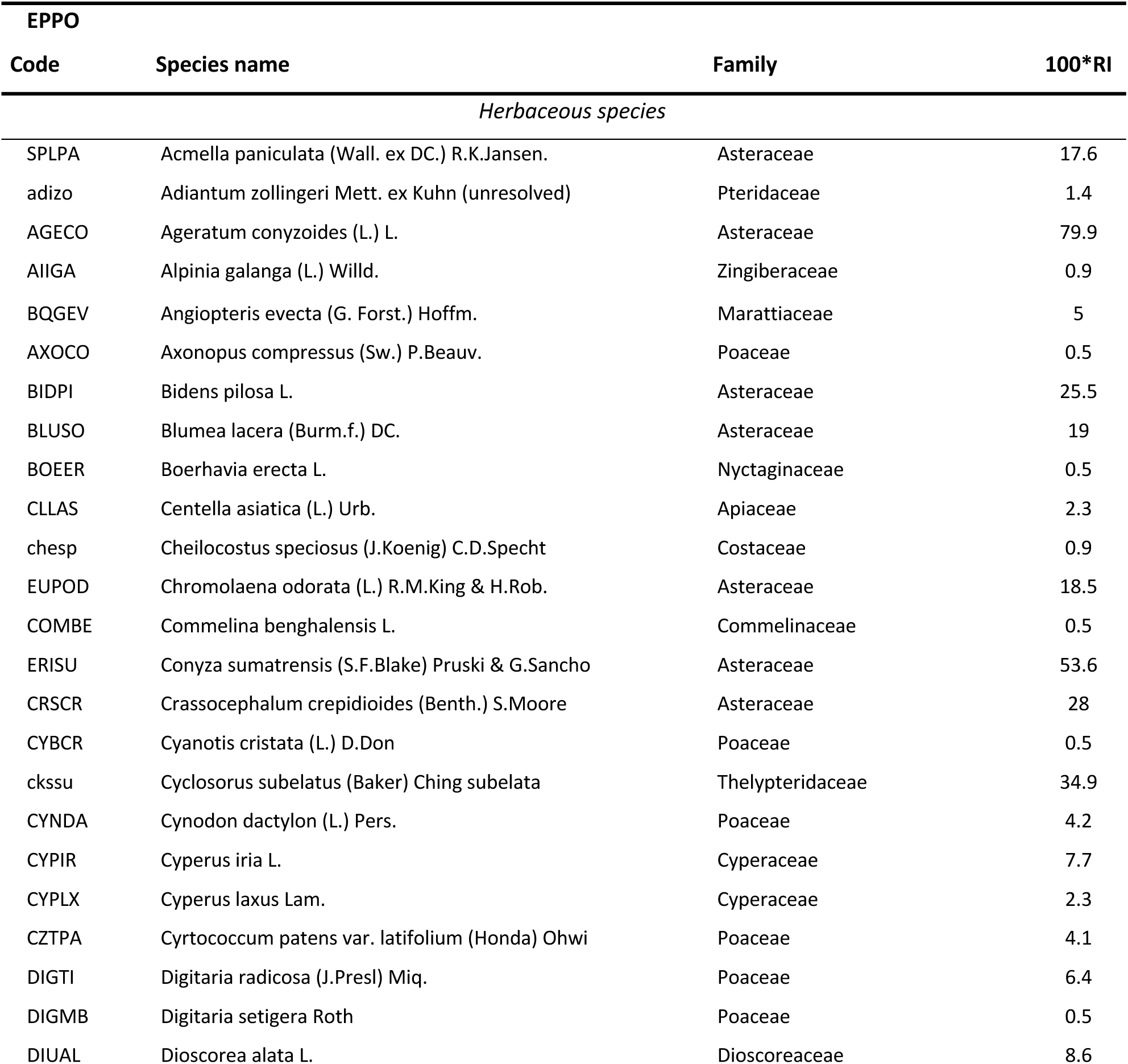
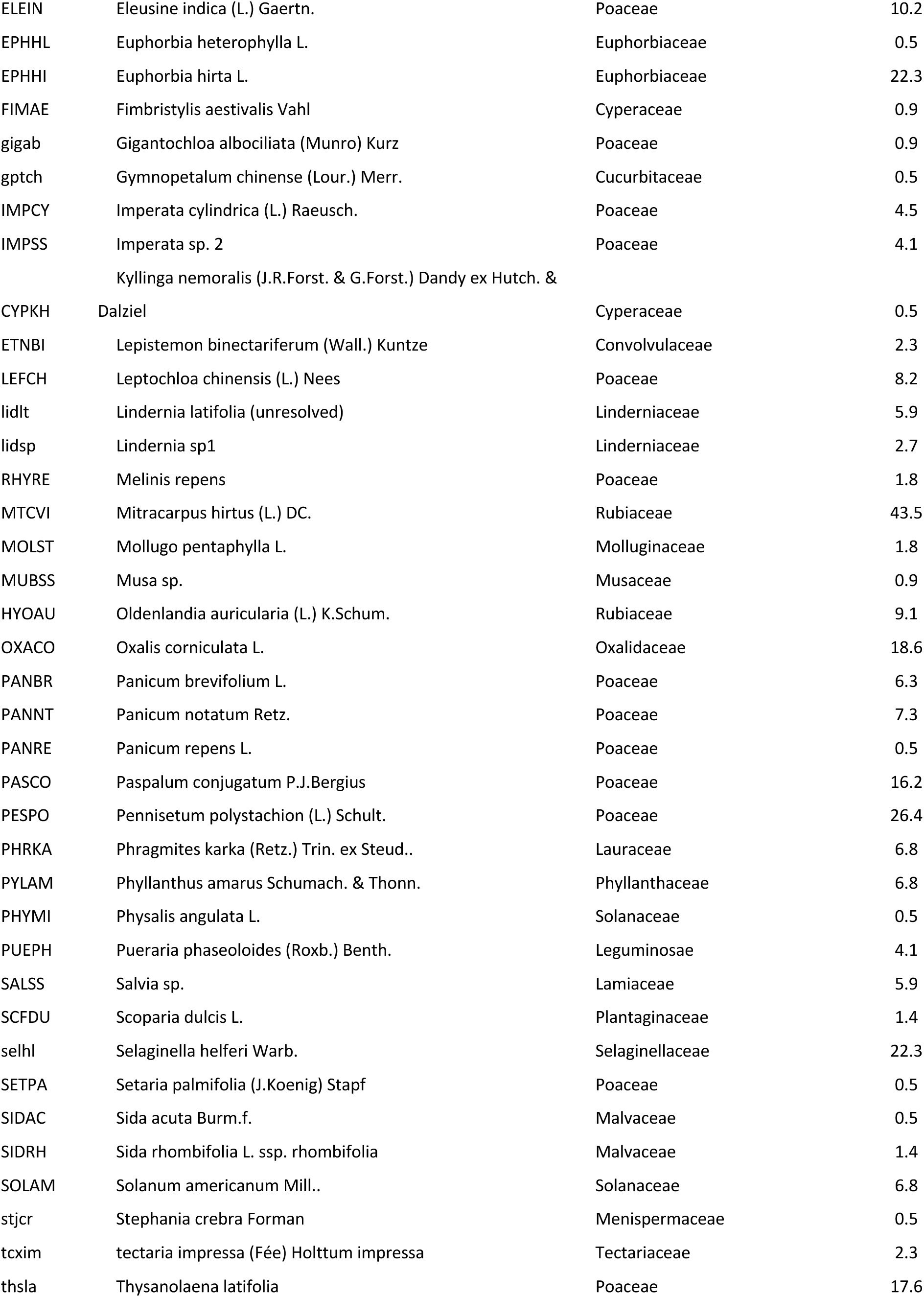
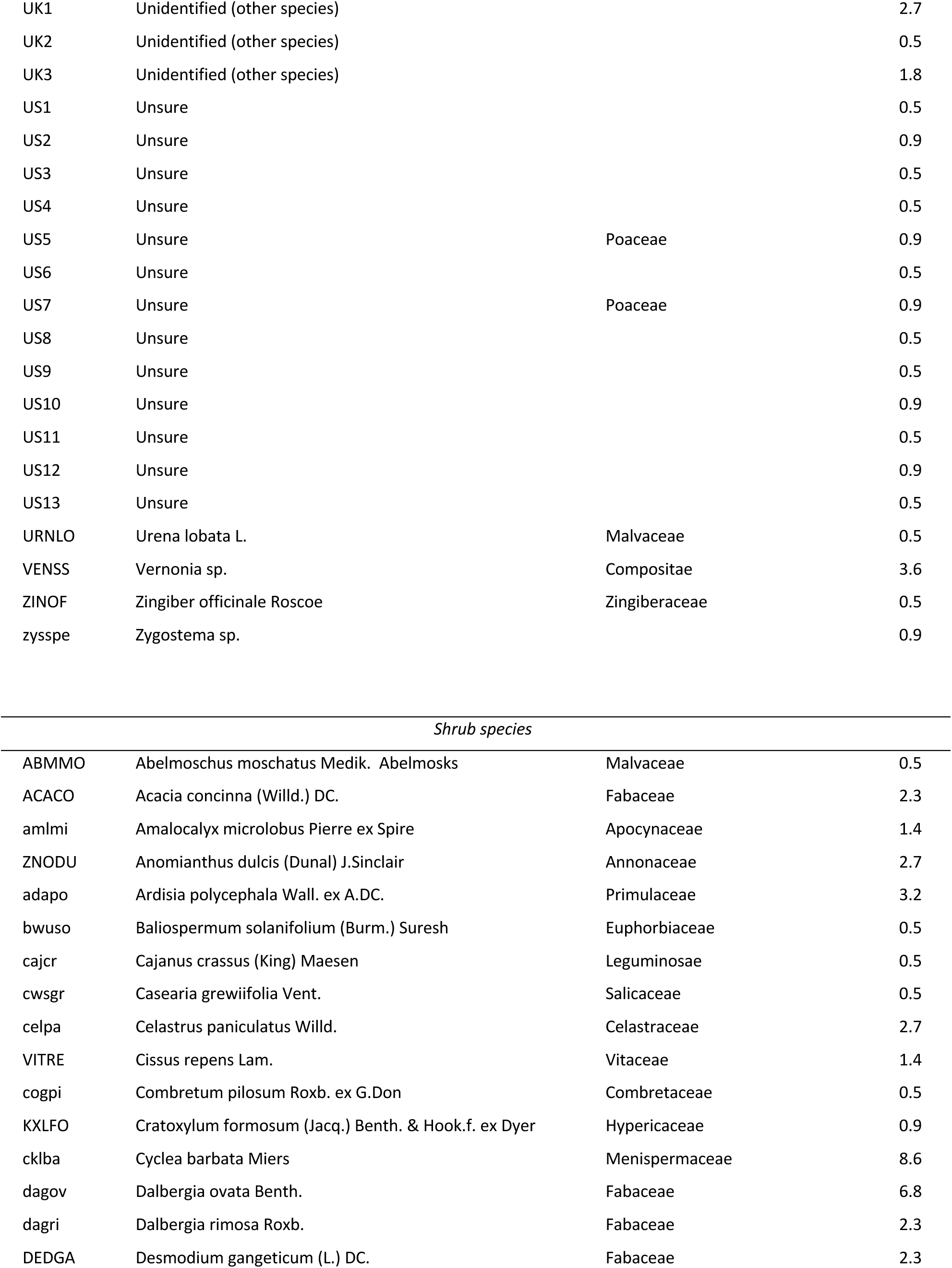
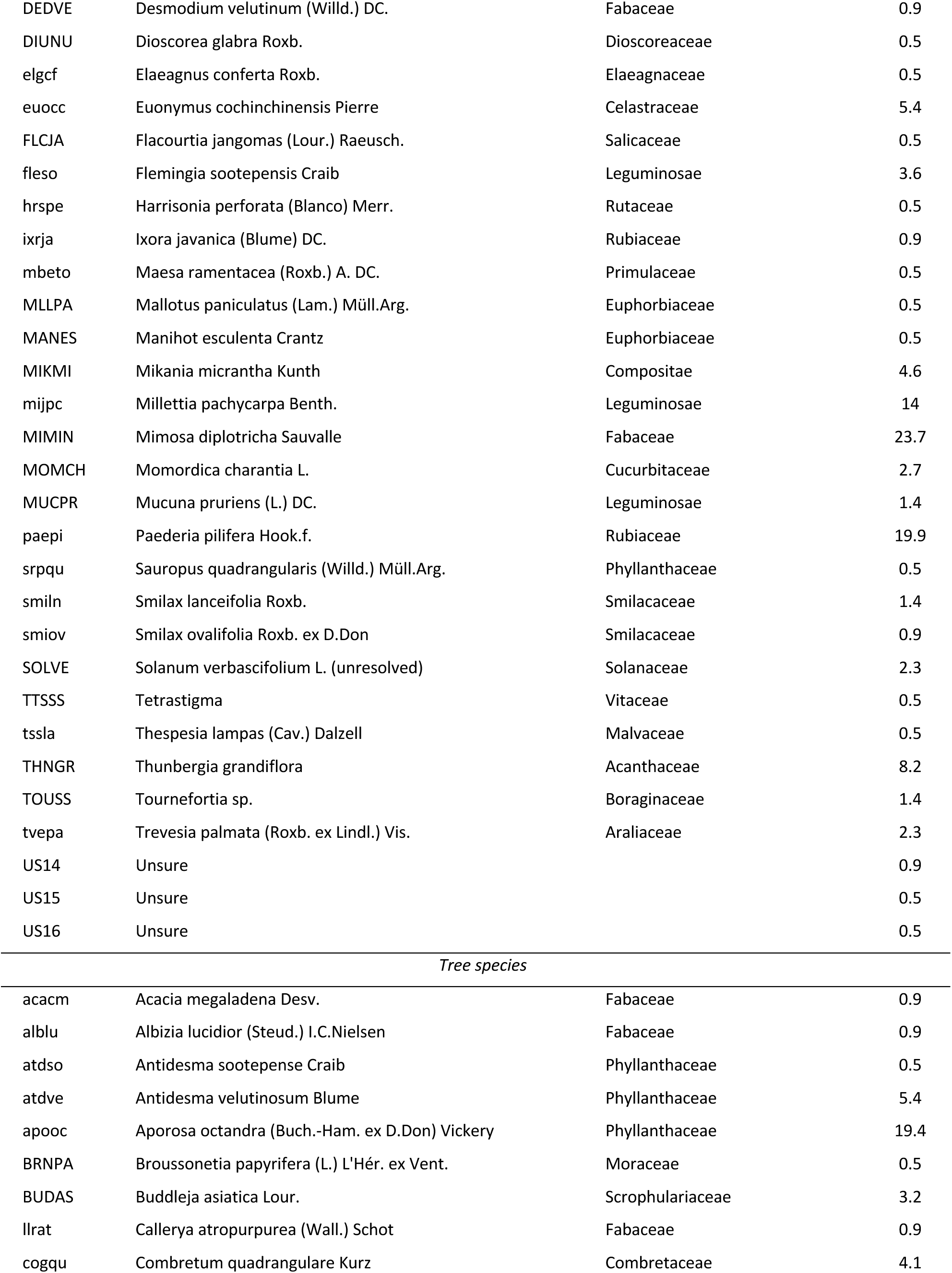
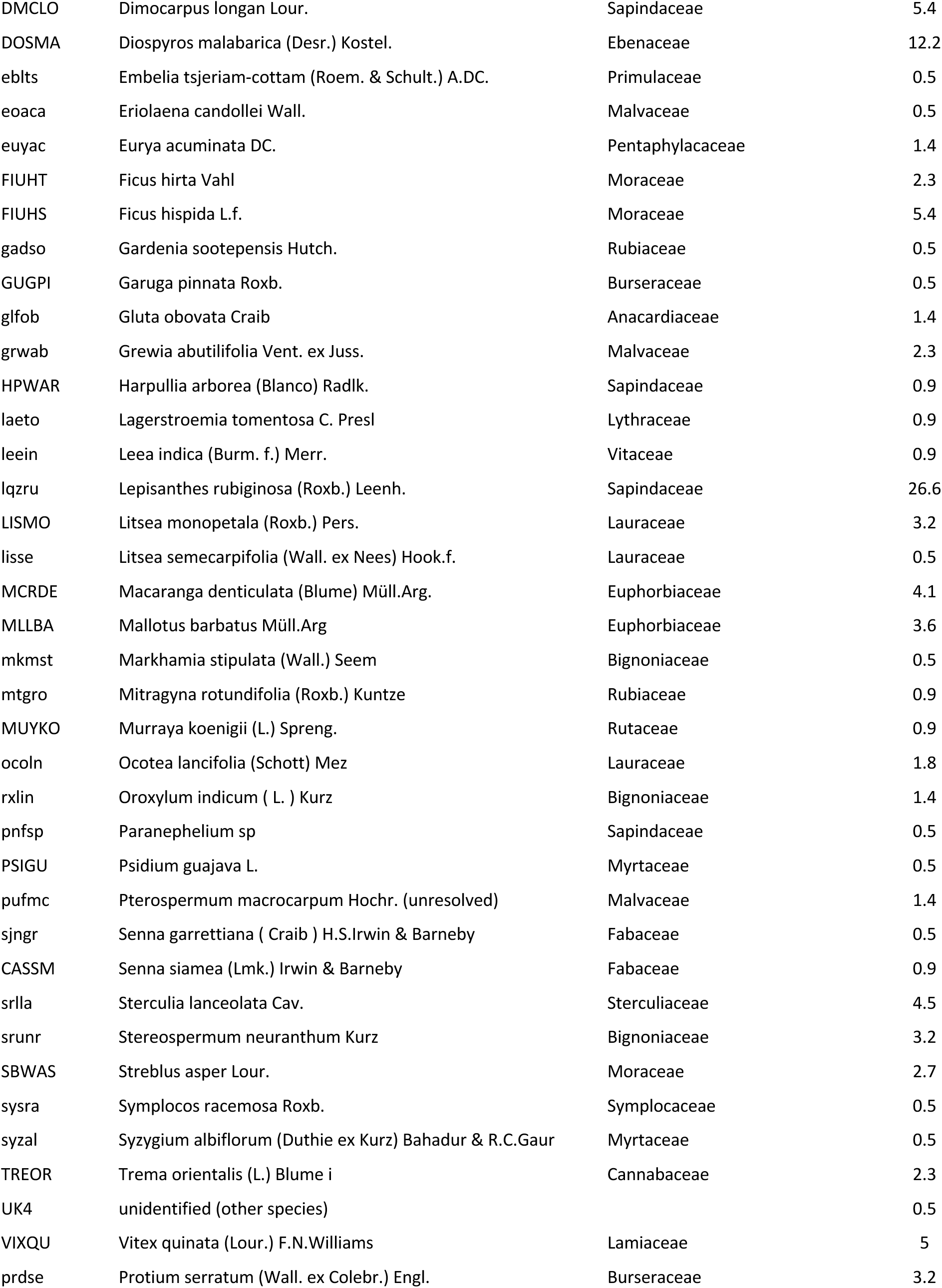
List of EPPO codes, plant species, and relative importance index multiplied by 100 (100*RI). RI (calculated as the mean of the frequency and relative abundance of the species in all plots) was multiplied by 100 to ease reading. EPPO codes in uppercase are official codes as per the EPPO database. Lowercase codes are custom codes for species absent from the database. Species labelled as “unidentified” correspond to species clearly different from the other species, but which we were unable to identify. Species labelled as “unsure” correspond to plants which might belong to one or the other of the identified species, but could not be identified with certainty (“unsure” species were not included in the analysis).

| EPPO |  |  |  |
| --- | --- | --- | --- |
| Code | Species name | Family | 100*RI |
| *Herbaceous species* |  |  |  |
| SPLPA | Acmella paniculata (Wall. ex DC.) R.K.Jansen. | Asteraceae | 17.6 |
| adizo | Adiantum zollingeri Mett. ex Kuhn (unresolved) | Pteridaceae | 1.4 |
| AGECO | Ageratum conyzoides (L.) L. | Asteraceae | 79.9 |
| AIIGA | Alpinia galanga (L.) Willd. | Zingiberaceae | 0.9 |
| BQGEV | Angiopteris evecta (G. Forst.) Hoffm. | Marattiaceae | 5 |
| AXOCO | Axonopus compressus (Sw.) P.Beauv. | Poaceae | 0.5 |
| BIDPI | Bidens pilosa L. | Asteraceae | 25.5 |
| BLUSO | Blumea lacera (Burm.f.) DC. | Asteraceae | 19 |
| BOEER | Boerhavia erecta L. | Nyctaginaceae | 0.5 |
| CLLAS | Centella asiatica (L.) Urb. | Apiaceae | 2.3 |
| chesp | Cheilocostus speciosus (J.Koenig) C.D.Specht | Costaceae | 0.9 |
| EUPOD | Chromolaena odorata (L.) R.M.King & H.Rob. | Asteraceae | 18.5 |
| COMBE | Commelina benghalensis L. | Commelinaceae | 0.5 |
| ERISU | Conyza sumatrensis (S.F.Blake) Pruski & G.Sancho | Asteraceae | 53.6 |
| CRSCR | Crassocephalum crepidioides (Benth.) S.Moore | Asteraceae | 28 |
| CYBCR | Cyanotis cristata (L.) D.Don | Poaceae | 0.5 |
| ckssu | Cyclosorus subelatus (Baker) Ching subelata | Thelypteridaceae | 34.9 |
| CYNDA | Cynodon dactylon (L.) Pers. | Poaceae | 4.2 |
| CYPIR | Cyperus iria L. | Cyperaceae | 7.7 |
| CYPLX | Cyperus laxus Lam. | Cyperaceae | 2.3 |
| CZTPA | Cyrtococcum patens var. latifolium (Honda) Ohwi | Poaceae | 4.1 |
| DIGTI | Digitaria radicata (J.Presl) Miq. | Poaceae | 6.4 |
| DIGMB | Digitaria setigera Roth | Poaceae | 0.5 |
| DIUAL | Dioscorea alata L. | Dioscoreaceae | 8.6 |
| ELEIN | Eleusine indica (L.) Gaertn. | Poaceae | 10.2 |
| EPHHL | Euphorbia heterophylla L. | Euphorbiaceae | 0.5 |
| EPHHI | Euphorbia hirta L. | Euphorbiaceae | 22.3 |
| FIMAE | Fimbristylis aestivalis Vahl | Cyperaceae | 0.9 |
| gigab | Gigantochloa albociliata (Munro) Kurz | Poaceae | 0.9 |
| gptch | Gymnopetalum chinense (Lour.) Merr. | Cucurbitaceae | 0.5 |
| IMPCY | Imperata cylindrica (L.) Raeusch. | Poaceae | 4.5 |
| IMPSS | Imperata sp. 2 | Poaceae | 4.1 |
|  | Kyllinga nemoralis (J.R.Forst. & G.Forst.) Dandy ex Hutch. & |  |  |
| CYPKH | Dalziel | Cyperaceae | 0.5 |
| ETNBI | Lepistemon binectariferum (Wall.) Kuntze | Convolvulaceae | 2.3 |
| LEFCH | Leptochloa chinensis (L.) Nees | Poaceae | 8.2 |
| lidlt | Lindernia latifolia (unresolved) | Linderniaceae | 5.9 |
| lidsp | Lindernia sp1 | Linderniaceae | 2.7 |
| RHYRE | Melinis repens | Poaceae | 1.8 |
| MTCVI | Mitracarpus hirtus (L.) DC. | Rubiaceae | 43.5 |
| MOLST | Mollugo pentaphylla L. | Molluginaceae | 1.8 |
| MUBSS | Musa sp. | Musaceae | 0.9 |
| HYOAU | Oldenlandia auricularia (L.) K.Schum. | Rubiaceae | 9.1 |
| OXACO | Oxalis corniculata L. | Oxalidaceae | 18.6 |
| PANBR | Panicum brevifolium L. | Poaceae | 6.3 |
| PANNT | Panicum notatum Retz. | Poaceae | 7.3 |
| PANRE | Panicum repens L. | Poaceae | 0.5 |
| PASCO | Paspalum conjugatum P.J.Bergius | Poaceae | 16.2 |
| PESPO | Pennisetum polystachion (L.) Schult. | Poaceae | 26.4 |
| PHRKA | Phragmites karka (Retz.) Trin. ex Steud.. | Lauraceae | 6.8 |
| PYLAM | Phyllanthus amarus Schumach. & Thonn. | Phyllanthaceae | 6.8 |
| PHYMI | Physalis angulata L. | Solanaceae | 0.5 |
| PUEPH | Pueraria phaseoloides (Roxb.) Benth. | Leguminosae | 4.1 |
| SALSS | Salvia sp. | Lamiaceae | 5.9 |
| SCFDU | Scoparia dulcis L. | Plantaginaceae | 1.4 |
| selhl | Selaginella helferi Warb. | Selaginellaceae | 22.3 |
| SETPA | Setaria palmifolia (J.Koenig) Stapf | Poaceae | 0.5 |
| SIDAC | Sida acuta Burm.f. | Malvaceae | 0.5 |
| SIDRH | Sida rhombifolia L. ssp. rhombifolia | Malvaceae | 1.4 |
| SOLAM | Solanum americanum Mill.. | Solanaceae | 6.8 |
| stjcr | Stephania crebra Forman | Menispermaceae | 0.5 |
| tcxim | tectaria impressa (Fée) Holttum impressa | Tectariaceae | 2.3 |
| thsla | Thysanolaena latifolia | Poaceae | 17.6 |
| UK1 | Unidentified (other species) |  | 2.7 |
| UK2 | Unidentified (other species) |  | 0.5 |
| UK3 | Unidentified (other species) |  | 1.8 |
| US1 | Unsure |  | 0.5 |
| US2 | Unsure |  | 0.9 |
| US3 | Unsure |  | 0.5 |
| US4 | Unsure |  | 0.5 |
| US5 | Unsure | Poaceae | 0.9 |
| US6 | Unsure |  | 0.5 |
| US7 | Unsure | Poaceae | 0.9 |
| US8 | Unsure |  | 0.5 |
| US9 | Unsure |  | 0.5 |
| US10 | Unsure |  | 0.9 |
| US11 | Unsure |  | 0.5 |
| US12 | Unsure |  | 0.9 |
| US13 | Unsure |  | 0.5 |
| URNLO | Urena lobata L. | Malvaceae | 0.5 |
| VENSS | Vernonia sp. | Compositae | 3.6 |
| ZINOF | Zingiber officinale Roscoe | Zingiberaceae | 0.5 |
| zysspe | Zygostema sp. |  | 0.9 |
| *Shrub species* |  |  |  |
| ABMMO | Abelmoschus moschatus Medik. Abelmosks | Malvaceae | 0.5 |
| ACACO | Acacia concinna (Willd.) DC. | Fabaceae | 2.3 |
| amlmi | Amalocalyx microlobus Pierre ex Spire | Apocynaceae | 1.4 |
| ZNODU | Anomianthus dulcis (Dunal) J.Sinclair | Annonaceae | 2.7 |
| adapo | Ardisia polycephala Wall. ex A.DC. | Primulaceae | 3.2 |
| bwuso | Baliospermum solanifolium (Burm.) Suresh | Euphorbiaceae | 0.5 |
| cajcr | Cajanus crassus (King) Maesen | Leguminosae | 0.5 |
| cwsgr | Casearia grewiifolia Vent. | Salicaceae | 0.5 |
| celpa | Celastrus paniculatus Willd. | Celastraceae | 2.7 |
| VITRE | Cissus repens Lam. | Vitaceae | 1.4 |
| cogpi | Combretum pilosum Roxb. ex G.Don | Combretaceae | 0.5 |
| KXLFO | Cratoxylum formosum (Jacq.) Benth. & Hook.f. ex Dyer | Hypericaceae | 0.9 |
| cklba | Cyclea barbata Miers | Menispermaceae | 8.6 |
| dagov | Dalbergia ovata Benth. | Fabaceae | 6.8 |
| dagri | Dalbergia rimosa Roxb. | Fabaceae | 2.3 |
| DEDGA | Desmodium gangeticum (L.) DC. | Fabaceae | 2.3 |
| DEDVE | Desmodium velutinum (Willd.) DC. | Fabaceae | 0.9 |
| DIUNU | Dioscorea glabra Roxb. | Dioscoreaceae | 0.5 |
| elgcf | Elaeagnus conferta Roxb. | Elaeagnaceae | 0.5 |
| euocc | Euonymus cochinchinensis Pierre | Celastraceae | 5.4 |
| FLCJA | Flacourtia jangomas (Lour.) Raeusch. | Salicaceae | 0.5 |
| fleso | Flemingia sootepensis Craib | Leguminosae | 3.6 |
| hrspe | Harrisonia perforata (Blanco) Merr. | Rutaceae | 0.5 |
| ixrja | Ixora javanica (Blume) DC. | Rubiaceae | 0.9 |
| mbeto | Maesa ramentacea (Roxb.) A. DC. | Primulaceae | 0.5 |
| MLLPA | Mallotus paniculatus (Lam.) Müll.Arg. | Euphorbiaceae | 0.5 |
| MANES | Manihot esculenta Crantz | Euphorbiaceae | 0.5 |
| MIKMI | Mikania micrantha Kunth | Compositae | 4.6 |
| mijpc | Milletia pachycarpa Benth. | Leguminosae | 14 |
| MIMIN | Mimosa diplotricha Sauvalle | Fabaceae | 23.7 |
| MOMCH | Momordica charantia L. | Cucurbitaceae | 2.7 |
| MUCPR | Mucuna pruriens (L.) DC. | Leguminosae | 1.4 |
| paepi | Paederia pilifera Hook.f. | Rubiaceae | 19.9 |
| srpqu | Sauropus quadrangularis (Willd.) Müll.Arg. | Phyllanthaceae | 0.5 |
| smiln | Smilax lanceifolia Roxb. | Smilacaceae | 1.4 |
| smiov | Smilax ovalifolia Roxb. ex D.Don | Smilacaceae | 0.9 |
| SOLVE | Solanum verbascifolium L. (unresolved) | Solanaceae | 2.3 |
| TTSSS | Tetrastigma | Vitaceae | 0.5 |
| tssla | Thespesia lampas (Cav.) Dalzell | Malvaceae | 0.5 |
| THNGR | Thunbergia grandiflora | Acanthaceae | 8.2 |
| TOUSS | Tournefortia sp. | Boraginaceae | 1.4 |
| tvepa | Trevesia palmata (Roxb. ex Lindl.) Vis. | Araliaceae | 2.3 |
| US14 | Unsure |  | 0.9 |
| US15 | Unsure |  | 0.5 |
| US16 | Unsure |  | 0.5 |
| *Tree species* |  |  |  |
| acacm | Acacia megaladena Desv. | Fabaceae | 0.9 |
| alblu | Albizia lucidior (Steud.) I.C.Nielsen | Fabaceae | 0.9 |
| atdso | Antidesma sootepense Craib | Phyllanthaceae | 0.5 |
| atdve | Antidesma velutinosum Blume | Phyllanthaceae | 5.4 |
| apooc | Aporosa octandra (Buch.-Ham. ex D.Don) Vickery | Phyllanthaceae | 19.4 |
| BRNPA | Broussonetia papyrifera (L.) L'Hér. ex Vent. | Moraceae | 0.5 |
| BUDAS | Buddleja asiatica Lour. | Scrophulariaceae | 3.2 |
| llrat | Callerya atropurpurea (Wall.) Schot | Fabaceae | 0.9 |
| cogqu | Combretum quadrangulare Kurz | Combretaceae | 4.1 |
| DMCLO | Dimocarpus longan Lour. | Sapindaceae | 5.4 |
| DOSMA | Diospyros malabarica (Desr.) Kostel. | Ebenaceae | 12.2 |
| ebllts | Embelia tsjeriam-cottam (Roem. & Schult.) A.DC. | Primulaceae | 0.5 |
| eoaca | Eriolaena candollei Wall. | Malvaceae | 0.5 |
| euyac | Eurya acuminata DC. | Pentaphylacaceae | 1.4 |
| FIUHT | Ficus hirta Vahl | Moraceae | 2.3 |
| FIUHS | Ficus hispida L.f. | Moraceae | 5.4 |
| gadso | Gardenia sootepensis Hutch. | Rubiaceae | 0.5 |
| GUGPI | Garuga pinnata Roxb. | Burseraceae | 0.5 |
| glfob | Gluta obovata Craib | Anacardiaceae | 1.4 |
| grwab | Grewia abutilifolia Vent. ex Juss. | Malvaceae | 2.3 |
| HPWAR | Harpullia arborea (Blanco) Radlk. | Sapindaceae | 0.9 |
| laeto | Lagerstroemia tomentosa C. Presl | Lythraceae | 0.9 |
| leein | Leea indica (Burm. f.) Merr. | Vitaceae | 0.9 |
| lqzru | Lepisanthes rubiginosa (Roxb.) Leenh. | Sapindaceae | 26.6 |
| LISMO | Litsea monopetala (Roxb.) Pers. | Lauraceae | 3.2 |
| lisce | Litsea semecarpifolia (Wall. ex Nees) Hook.f. | Lauraceae | 0.5 |
| MCRDE | Macaranga denticulata (Blume) Müll.Arg. | Euphorbiaceae | 4.1 |
| MLLBA | Mallotus barbatus Müll.Arg | Euphorbiaceae | 3.6 |
| mkmst | Markhamia stipulata (Wall.) Seem | Bignoniaceae | 0.5 |
| mtgro | Mitragyna rotundifolia (Roxb.) Kuntze | Rubiaceae | 0.9 |
| MUYKO | Murraya koenigii (L.) Spreng. | Rutaceae | 0.9 |
| ocoln | Ocotea lancifolia (Schott) Mez | Lauraceae | 1.8 |
| rxlin | Oroxylum indicum ( L. ) Kurz | Bignoniaceae | 1.4 |
| pnfsp | Paranephelium sp | Sapindaceae | 0.5 |
| PSIGU | Psidium guajava L. | Myrtaceae | 0.5 |
| pufmc | Pterospermum macrocarpum Hochr. (unresolved) | Malvaceae | 1.4 |
| sjngr | Senna garrettiana ( Craib ) H.S.Irwin & Barneby | Fabaceae | 0.5 |
| CASSM | Senna siamea (Lmk.) Irwin & Barneby | Fabaceae | 0.9 |
| srlla | Sterculia lanceolata Cav. | Sterculiaceae | 4.5 |
| srntr | Stereospermum neuranthum Kurz | Bignoniaceae | 3.2 |
| SBWAS | Streblus asper Lour. | Moraceae | 2.7 |
| sysra | Symplocos racemosa Roxb. | Symplocaceae | 0.5 |
| syzal | Syzygium albiflorum (Duthie ex Kurz) Bahadur & R.C.Gaur | Myrtaceae | 0.5 |
| TREOR | Trema orientalis (L.) Blume i | Cannabaceae | 2.3 |
| UK4 | unidentified (other species) |  | 0.5 |
| VIXQU | Vitex quinata (Lour.) F.N.Williams | Lamiaceae | 5 |
| prdse | Protium serratum (Wall. ex Colebr.) Engl. | Burseraceae | 3.2 |

**Table S2.** Effects of crop type and number of crop types on weed richness, diversity and abundance: results from mixed model analyses.

|  | Number<br>of crop<br>types | Seaso<br>n | Crop types<br>x season | Annual<br>crop | Presenc<br>e of trees | Global<br>R<sup>2</sup> |
| --- | --- | --- | --- | --- | --- | --- |
| Richness (all species) | 4 n.s | 1 n.s | 6 n.s | 0 n.s | 0 n.s | 15 |
| Richness (herbaceous species) | 10 * | 1 n.s | 1 n.s | 4 n.s | 3 n.s | 37 |
| Richness (tree and shrub species) | 1 n.s | 1 n.s | 11 . | 2 n.s | 3 n.s | 21 |
| Dominance (all species) | 9 * | 17 *** | 0 n.s | 9 * | 0 n.s | 27 |
| Dominance (herbaceous species) | 11 ** | 13 ** | 2 n.s | 12 ** | 1 n.s | 28 |
| Dominance (tree shrub species) | 0 n.s | 1 n.s | 0 n.s | 3 n.s | 2 n.s | 5 |
| Shannon index (all species) | 5 . | 14 *** | 2 n.s | 2 n.s | 0 n.s | 20 |
| Shannon (herbaceous species) | 6 * | 10 ** | 6 n.s | 3 n.s | 3 n.s | 25 |
| Shannon (tree and shrub species) | 0 n.s | 0 n.s | 0 n.s | 2 n.s | 1 n.s | 4 |
| Biomass (square-root transformed, all species) | 3 n.s | 4 n.s | 4 n.s | 8 . | 9 * | 20 |
| Density (log, all species) | 0 n.s | 11 * | 1 n.s | 1 n.s | 1 n.s | 15 |

|  | Number<br>of crop<br>types | Estimate (dry season) | Estimate (rainy season) |
| --- | --- | --- | --- |
| Species number (all species) | 1 | 15.6 (11.4 - 19.9) <sup>a</sup> | 13.4 (8 - 18.8) <sup>a</sup> |
|  | 2 | 16.5 (13.9 - 19) <sup>a</sup> | 19.3 (16.6 - 22) <sup>a</sup> |
|  | 3 | 16.5 (11.8 - 21.1) <sup>a</sup> | 18.8 (13.8 - 23.8) <sup>a</sup> |
| Species number (herbaceous species) | 1 | 8.2 (5.4 - 11) <sup>a</sup> | 8.8 (5.1 - 12.5) <sup>a</sup> |
|  | 2 | 11.7 (9.9 - 13.4) <sup>a</sup> | 12.7 (10.8 - 14.6) <sup>a</sup> |
|  | 3 | 12.3 (9.2 - 15.5) <sup>a</sup> | 12.8 (9.5 - 16.2) <sup>a</sup> |
| Species number (tree and shrub species) | 1 | 7.5 (4.5 - 10.5) <sup>a</sup> | 4.9 (1.1 - 8.7) <sup>a</sup> |
|  | 2 | 4.9 (3.1 - 6.7) <sup>a</sup> | 6.6 (4.7 - 8.5) <sup>a</sup> |
|  | 3 | 3.8 (0.5 - 7) <sup>a</sup> | 6.2 (2.7 - 9.7) <sup>a</sup> |
| **Dominance (all species)** | 1 | **0.8 (0.6 - 0.9) <sup>ab</sup>** | 0.9 (0.7 - 1.1) <sup>a</sup> |
|  | 2 | **0.5 (0.4 - 0.6) <sup>a</sup>** | 0.7 (0.6 - 0.8) <sup>a</sup> |
|  | 3 | **0.5 (0.3 - 0.7) <sup>b</sup>** | 0.7 (0.5 - 0.9) <sup>a</sup> |
| **Dominance (herbaceous species)** | 1 | **0.8 (0.7 - 1) <sup>a</sup>** | 0.9 (0.7 - 1.1) <sup>a</sup> |
|  | 2 | **0.6 (0.5 - 0.7) <sup>a</sup>** | 0.7 (0.6 - 0.8) <sup>a</sup> |
|  | 3 | **0.6 (0.4 - 0.7) <sup>b</sup>** | 0.7 (0.6 - 0.9) <sup>a</sup> |
| Dominance | 1 | 0.7 (0.5 - 0.9) <sup>a</sup> | 0.7 (0.5 - 1) <sup>a</sup> |
| (tree and shrub species) | 2 | 0.7 (0.6 - 0.8) <sup>a</sup> | 0.7 (0.6 - 0.8) <sup>a</sup> |
|  | 3 | 0.7 (0.5 - 0.9) <sup>a</sup> | 0.7 (0.5 - 1) <sup>a</sup> |
| **Shannon index** | 1 | **0.9 (0.6 - 1.2) <sup>a</sup>** | 0.7 (0.3 - 1.2) <sup>a</sup> |
| **(all species)** | 2 | **1.3 (1.1 - 1.5) <sup>ab</sup>** | 1 (0.7 - 1.2) <sup>a</sup> |
|  | 3 | **1.5 (1.1 - 1.9) <sup>b</sup>** | 0.9 (0.5 - 1.3) <sup>a</sup> |
| **Shannon** | 1 | **0.6 (0.3 - 1) <sup>a</sup>** | 0.6 (0.2 - 1) <sup>a</sup> |
| **(herbaceous species)** | 2 | **1 (0.9 - 1.2) <sup>ab</sup>** | 0.8 (0.6 - 1) <sup>a</sup> |
|  | 3 | **1.3 (1 - 1.7) <sup>b</sup>** | 0.8 (0.4 - 1.1) <sup>a</sup> |
| Shannon | 1 | 0.7 (0.3 - 1.1) <sup>a</sup> | 0.7 (0.1 - 1.2) <sup>a</sup> |
| (tree and shrub species) | 2 | 0.7 (0.4 - 1) <sup>a</sup> | 0.6 (0.4 - 0.9) <sup>a</sup> |
|  | 3 | 0.7 (0.2 - 1.1) <sup>a</sup> | 0.7 (0.2 - 1.2) <sup>a</sup> |
| Biomass | 1 | 7.8 (4 - 11.7) <sup>a</sup> | 3.3 (-1.8 - 8.5) <sup>a</sup> |
| (square-root | 2 | 8.8 (6.6 - 11.1) <sup>a</sup> | 7.8 (5.3 - 10.2) <sup>a</sup> |
| transformed, all species) | 3 | 7.9 (3.5 - 12.2) <sup>a</sup> | 7.9 (3.2 - 12.6) <sup>a</sup> |
| Density | 1 | 5.3 (4.4 - 6.2) <sup>a</sup> | 6.1 (4.9 - 7.2) <sup>a</sup> |
| (log-transformed, all | 2 | 5.5 (5 - 6.1) <sup>a</sup> | 6 (5.4 - 6.5) <sup>a</sup> |
| species) | 3 | 5.4 (4.4 - 6.4) <sup>a</sup> | 6.2 (5.2 - 7.3) <sup>a</sup> |

**Table S3.** Effects of the presence of trees and crop shifts on weed richness, diversity and abundance: results from mixed model analyses including only fields with maize as the current annual crop.

|  | Number of<br>crop types | Season | Crop types<br>x season | Presence<br>of trees | Global R<sup>2</sup> |
| --- | --- | --- | --- | --- | --- |
| Species number (all species) | 30 * | 6 n.s | 33 ** | 6 . | 37 |
| Species number (herbaceous species) | 50 *** | 13 n.s | 11 n.s | 1 n.s | **41** |
| Species number (tree and shrub species) | 0 n.s | 0 n.s | 27 ** | 14 ** | 34 |
| Dominance (all species) | 18 * | 75 *** | 19 . | 24 ** | 71 |
| Dominance (herbaceous species) | 19 . | 74 *** | 27 * | 30 *** | 72 |
| Dominance (tree shrub species) | 3 n.s | 3 n.s | 2 n.s | 3 n.s | 8 |
| Shannon index (all species) | 27 ** | 59 *** | 23 * | 11 . | 59 |
| Shannon (herbaceous species) | 17 . | 55 ** | 33 * | 25 ** | 59 |
| Shannon (tree and shrub species) | 1 n.s | 2 n.s | 1 n.s | 4 n.s | 7 |
| Biomass (square-root transformed, all species) | 54 * | 7 * | 24 ** | 23 ** | 51 |
| Density (log, all species) | 47 * | 48 *** | 14 n.s | 0 n.s | 58 |

|  | Number of<br>crop shifts | Estimate (dry season) | Estimate (rainy season) |
| --- | --- | --- | --- |
| **Species number (all species)** | 0 | 14.8 (10.8 - 18.9) <sup>a</sup> | **11.9 (6.4 - 17.4) <sup>a</sup>** |
|  | 2 | 17.7 (10.5 - 24.9) <sup>a</sup> | **24.3 (16.8 - 31.8) <sup>b</sup>** |
| **Species number (herbaceous species)** | 0 | **7.4 (4.6 - 10.2) <sup>a</sup>** | **7.5 (3.5 - 11.4) <sup>a</sup>** |
|  | 2 | **13 (8.1 - 17.9) <sup>b</sup>** | **16.3 (11.1 - 21.5) <sup>b</sup>** |
| Species number (tree and shrub species) | 0 | 7.5 (4.7 - 10.4) <sup>a</sup> | 4.6 (0.6 - 8.5) <sup>a</sup> |
|  | 2 | 4.7 (-0.3 - 9.8) <sup>a</sup> | 7.8 (2.5 - 13.2) <sup>a</sup> |
| Dominance (all species) | 0 | 0.6 (0.5 - 0.7) <sup>a</sup> | 0.8 (0.7 - 0.9) <sup>a</sup> |
|  | 2 | 0.5 (0.3 - 0.6) <sup>a</sup> | 0.8 (0.6 - 0.9) <sup>a</sup> |
| Dominance (herbaceous species) | 0 | 0.7 (0.6 - 0.8) <sup>a</sup> | 0.8 (0.7 - 1) <sup>a</sup> |
|  | 2 | 0.5 (0.3 - 0.7) <sup>a</sup> | 0.7 (0.5 - 0.9) <sup>a</sup> |
| Dominance (tree and shrub species) | 0 | 0.7 (0.5 - 0.9) <sup>a</sup> | 0.7 (0.5 - 0.9) <sup>a</sup> |
|  | 2 | 0.7 (0.4 - 1) <sup>a</sup> | 0.6 (0.3 - 0.9) <sup>a</sup> |
| **Shannon index (all species)** | 0 | **1.1 (0.8 - 1.3) <sup>a</sup>** | 0.8 (0.4 - 1.1) <sup>a</sup> |
|  | 2 | **1.6 (1.3 - 2) <sup>b</sup>** | 0.8 (0.4 - 1.2) <sup>a</sup> |
| Shannon (herbaceous species) | 0 | 0.8 (0.5 - 1.1) <sup>a</sup> | 0.7 (0.3 - 1) <sup>a</sup> |
|  | 2 | 1.3 (0.8 - 1.8) <sup>a</sup> | 0.9 (0.3 - 1.5) <sup>a</sup> |
| Shannon | 0 | 0.7 (0.3 - 1.1) <sup>a</sup> | 0.7 (0.2 - 1.3) <sup>a</sup> |
| (tree and shrub species) | 2 | 0.7 (0 - 1.5) <sup>a</sup> | 0.9 (0.2 - 1.7) <sup>a</sup> |
| **Biomass** | 0 | 6.3 (4.1 - 8.5) <sup>a</sup> | **1.5 (-2.1 - 5) <sup>a</sup>** |
| **(square-root** | 2 | 7.3 (3.4 - 11.1) <sup>a</sup> | **9.2 (5 - 13.4) <sup>b</sup>** |
| **transformed, all species)** |  |  |  |
| **Density** | 0 | 5.2 (4.6 - 5.7) <sup>a</sup> | **5.9 (5 - 6.8) <sup>a</sup>** |
| **(log-transformed, all** | 2 | 5.6 (4.7 - 6.5) <sup>a</sup> | **7.4 (6.4 - 8.4) <sup>b</sup>** |
| **species)** |  |  |  |

**Figure S1.**
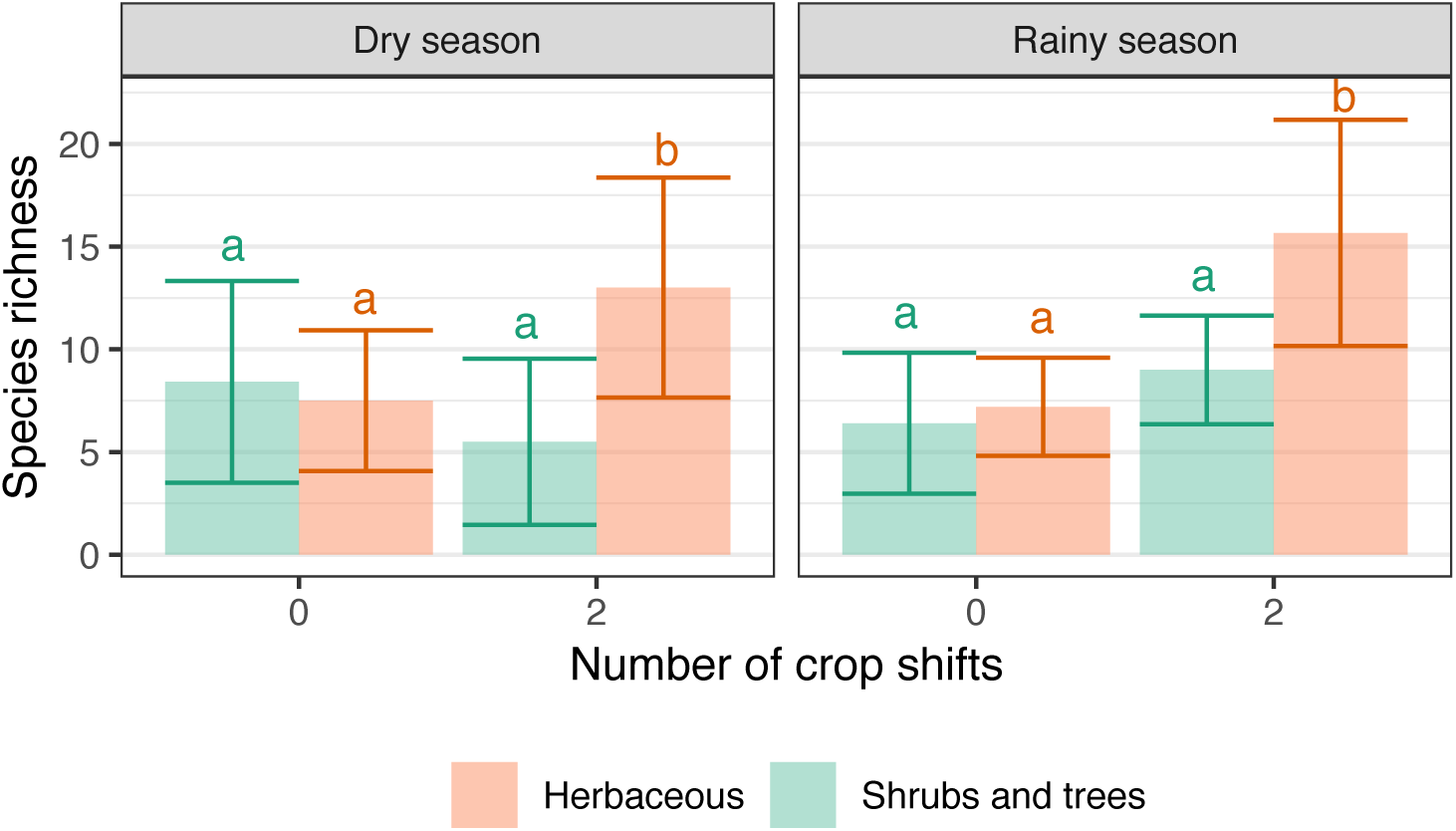
Variation of species richness per field with the number of crop shifts for fields whose current annual crop is maize. Bars represent the mean + /- standard deviation. Different letters indicate significant differences within each group (P < 0.05).

